# Glycation Stress–Driven Transcriptomic Signature Predicts Survival Benefit from Adjuvant Gemcitabine in Resectable Pancreatic Cancer

**DOI:** 10.1101/2025.07.07.663493

**Authors:** Oier Azurmendi Senar, Kosta Stosic, Christelle Bouchart, Julie Navez, Jawad Tarfouss, Rebekah Crake, Laurine Verset, Pieter Demetter, Marjorie Mauduit, Thierry Conroy, Jérôme Cros, Remy Nicolle, Manuel Saiselet, Joel Rodrigues Vitoria, Vincent Detours, Akeila Bellahcène, Tatjana Arsenijevic, Jean-Luc Van-Laethem, PRODIGE-24/CCTG PA6 consortium, MOSAPAC consortium

**Author notes:** |aCORRESPONDING AUTHOR Jean-Luc Van Laethem, Laboratory of Experimental Gastroenterology, Université Libre de Bruxelles, Route de Lennik 808, 1070 Brussels, Belgium,. Authors contributed equally (Akeila Bellahcène, Tatjana Arsenijevic, Jean-Luc Van-Laethem).

## Abstract

**BACKGROUND & AIMS:** Pancreatic ductal adenocarcinoma (PDAC) is a highly lethal cancer with limited response to systemic therapy. Gemcitabine (GEM) benefits only a subset of patients. Methylglyoxal (MG), a glycolysis byproduct, has been linked to tumor behavior and therapy response in PDAC, suggesting potential as a stratification marker.

**METHODS:** We developed a metabolically informed gene signature (MG-GEM) integrating MG-related glycolytic stress with clinical outcomes. Using the Puleo cohort (n=309), differential expression analysis between tumors with high and low MG stress identified 365 genes. LASSO Cox regression selected 16 prognostic genes, combined into a weighted risk score for patient stratification. MG GEM was validated in internal and external cohorts, including PRODIGE 24/CCTG PA6. Molecular, transcriptomic, and immune features were compared between high and low MG GEM groups. Finally, predictive performance was evaluated against the GemPred signature.

**RESULTS:** MG GEM divided PDAC patients into distinct risk groups with marked differences in overall and disease-free survival among GEM treated patients (OS 11.7 vs 27.2 months; DFS 7.6 vs 17.8 months, both p<0.0001). High MG-GEM tumors showed enrichment for KRAS G12D and SMAD4 mutations, basal and activated stroma subtypes, glycolytic metabolism, and reduced immune infiltration. Low MG-GEM tumors showed KRAS G12V, classical and immune subtypes, cholesterogenic metabolism, and adaptive immune favourable signatures. MG-GEM independently predicted GEM-specific clinical outcomes, irrespective of GemPred signature, and significantly enhanced patient stratification when combined with it. Within the PRODIGE-24/TGCC PA6 cohort, MG-GEM exhibited prognostic relevance and selectively identified patients who derived a survival benefit from adjuvant GEM, but not from FOLFIRINOX.

**CONCLUSIONS:** The 16 gene MG GEM signature predicts prognosis in resected PDAC, reflects glycolytic stress driven chemoresistance, surpassing conventional molecular classifications. As a metabolically informed signature, MG-GEM holds promise for guiding chemotherapy selection and informing KRAS-targeted combination strategies, meriting further prospective clinical validation.

## Introduction

Pancreatic ductal adenocarcinoma (PDAC), projected to become the second leading cause of cancer-related death by 2030 [1], is highly aggressive, with a 5-year overall survival (OS) rate below 12% [2]. Progress in developing more effective chemotherapy (CT) regimens has been limited [3], [4], [5] and clinical trials testing immunotherapy or targeted agents have, so far, failed to change the current clinical practice [6], [7] Currently, two CT regimens form the backbone of PDAC treatment: full-dose or modified FOLFIRINOX (folinic acid, 5-fluorouracil, irinotecan, and oxaliplatin; mFFX) and gemcitabine combined with nab-paclitaxel (GEM/nab-P) [8]. Their efficacy is broadly comparable, and choice between them is primarily guided by clinical criteria.

Gemcitabine (GEM) remains a cornerstone of systemic therapy for PDAC, yet only a subset of patients derives meaningful benefit. To address this challenge, transcriptomic classifiers have been developed to stratify patients based on their likelihood of responding to GEM. One such signature, GemPred [9], derived from preclinical models including cell lines and organoids, identifies approximately 26% of PDAC patients as predicted responders to GEM-based regimens. Additional transcriptomic tools have also been proposed to guide therapeutic decision-making. For example, GemCore [10], an integrative signature combining four independent GEM-response classifiers, identifies ∼45% of patients as GEM-sensitive. Similarly, Pancreas-View [11], which integrates transcriptomic data from preclinical models with machine learning to provide PDAC molecular subtyping and therapy-response predictions, identifies ∼37% of patients as GEM-sensitive. Together, these efforts underscore the ongoing commitment to optimizing precision treatment for PDAC.

In recent years, accumulating evidence has underscored the critical role of reprogrammed metabolism in shaping cancer treatment response and prognosis. In pancreatic cancer cells, genetic mutations are notably implicated in driving distinct metabolic phenotypes, including the Warburg effect [12]. Through this altered metabolic state, cancer cells increase glucose uptake and preferentially ferment glucose to lactate, even in the presence of oxygen [13]. Enhanced glycolytic activity inevitably results in the production of methylglyoxal (MG), a highly reactive aldehyde. Its accumulation and the formation of associated protein adduct, observed at elevated levels in breast, colon, and pancreatic cancers [14], underscore its significant role in tumor progression and chemoresistance.

In PDAC, previous studies have also linked glycolytic metabolism to chemoresistance and poor clinical outcomes [15], [16]. Aberrant KRAS activation, a hallmark of PDAC, drives enhanced glucose uptake and glycolysis through the upregulation of GLUT1 and key metabolic enzymes [12], [17]. A recent mechanistic study demonstrated that increased glycolytic flux in PDAC models supports de novo pyrimidine biosynthesis, thereby reducing GEM efficacy through a competitive metabolic mechanism [18].Interestingly, GEM-resistant cancer cells display elevated levels of glycolytic intermediates upstream of dihydroxyacetone phosphate and glyceraldehyde-3- phosphate [18], both of which serve as precursors for the spontaneous formation of MG [19]. Moreover, we have shown in PDAC cell models that GEM treatment itself induces accumulation of MG-adducts, and resistant cells adapt to MG stress by upregulating GLO1, the key MG-detoxifying enzyme [20]. Together, these findings highlight the biological relevance of MG stress in GEM resistance and support a metabolism-focused scoring strategy centred on MG, which we found to correlate with poorer overall survival in GEM-treated PDAC patients [20]. To operationalize this, MG stress levels in chemotherapy-naïve resected PDAC tumors were assessed using two complementary indicators: expression of GLO1 and of a panel of ten glycolysis-related genes. This approach enabled the classification of PDAC tumors into three MG stress categories: high MG stress, low MG stress, and Intermediate. Although the MG stress score was prognostic, its utility was limited to a small subset of the cohort we explored, as majority of tumors fell into the intermediate category [20].

This study aimed to refine the MG-stress score into a prognostically optimized gene signature that integrates MG stress–associated biology with prediction of clinical outcome in GEM receiving resectable pancreatic cancer. To achieve this, we applied LASSO Cox regression to 365 differentially expressed genes between high- and low-MG stress tumors, identifying a subset of genes most strongly associated with prognosis. These genes were combined into a weighted risk score, forming the 16- gene MG-GEM signature, which enables patient stratification based on predicted clinical outcomes.

We further explored the clinical and therapeutic relevance of the MG-GEM signature in both internal and external cohorts, demonstrating its predictive value as a tool for selecting PDAC patients likely to benefit from adjuvant GEM treatment.

## Materials and Methods

### Patient cohorts and study design

This study analyzed three cohorts of patients with pancreatic ductal adenocarcinoma (PDAC) using the following criteria: inclusion of patients ≥18 years old with complete clinicopathological data and no preoperative metastatic disease; exclusion of non-ductal histology and death within 30 days of surgery.

1. Puleo et al. cohort (n = 309, 1999–2010): Treatment-naïve, surgically resected PDAC cases with comprehensive clinical annotations and microarray-based transcriptomic data. The 57.28% (n=177) of patients of the cohort received adjuvant GEM, 31.71% (n=98) of the patients did not receive any adjuvant treatment (No AT) and the 7.76% (n=24) received other adjuvant strategies (Other AT). A subset of 89 patients (GEM: 64.05% (n=57); No AT: 28.09% (n=25); Other AT: 4.5% (n=4)) served as the discovery set for the derivation of the MG-GEM signature [21], and 220 patients (GEM: 54.55% (n=120); No AT: 33.18% (n=73); Other AT: 9.09% (n=20)) were used for the signature validation (Figure 1-A).
2. Erasme University Hospital cohort (n = 60, 2011–2020): Retrospective cohort of treatment-naïve, surgically resected PDAC cases with available FFPE tumor specimens for transcriptome profiling [22]. Most patients received adjuvant chemotherapy, and detailed follow-up allowed assessment of the prognostic and predictive value of the MG-GEM signature.
3. PRODIGE24/CCTG PA6 phase III trial cohort (n = 350; EudraCT: 2011-002026-52): Randomized trial comparing modified FOLFIRINOX versus GEM as adjuvant therapy [23], providing an external validation dataset to evaluate the predictive performance of MG-GEM for GEM survival relative to FOLFIRINOX.

**Figure 1.**
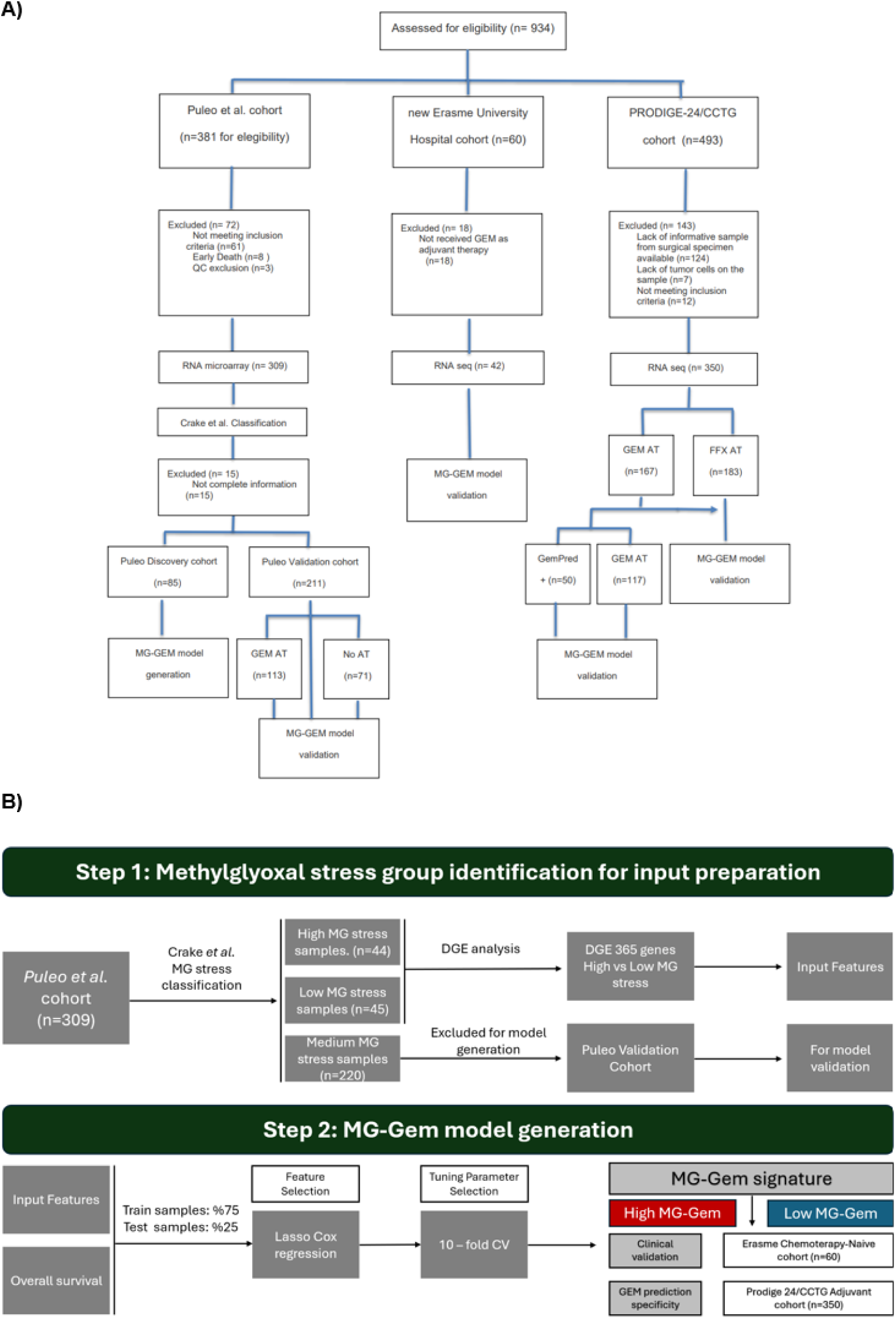
Workflow of the study. **(A)** CONSORT diagram of the procedure followed in the work. **(B)** Overview of the study workflow. **Abbreviations:** MG, Methylglyoxal; DGE, Differential Gene Expression; CV, Cross- Validation; GEM, Gemcitabine.

Figure 1-A shows a CONSORT-style flow diagram detailing patient cohorts used in this study for MG-GEM model generation and validation.

### Data acquisition and processing

The discovery cohort consisted of 309 samples from the Puleo et al. study [21], profiled on Affymetrix HGU219 RNA microarrays, with probe-level intensities processed using Robust Multi-array Average (RMA) normalization. For the PRODIGE-24/CCTG PA6 dataset [3], raw gene counts were normalized via the Upper Quartile method and subsequently log₂-transformed prior to downstream analyses.

### Transcriptomic profile process and analysis

Patients from the Puleo cohort (n=309) [21] were classified into high (n=44) and low (n=45) MG stress tumors using the method of Crake et al. [20] (Figure 1-B). This classification combines the expression of 10 glycolytic genes with that of GLO1, the main MG-detoxifying enzyme. Differential gene expression (DGE) analysis was performed between patients with different MG-stress status using the R-packages edgeR v3.40.2 and limma v3.54.2 packages [24], [25]. Genes were considered differentially expressed when log two-fold change was at least ±1 and p-adjusted value < 0.05. The Pancreatic Adenocarcinoma Molecular Gradient (PAMG) classifier was applied to determine the aggressiveness of the samples [26].

### Construction of the MG-GEM signature

Samples used for DGE analysis were randomly separated into a training set (75%) and a testing set (25%). Least Absolute Shrinkage and Selection Operator (LASSO) penalized Cox regression was conducted using the glmnet v4.1-8 package [27]. The RMA-normalized expression values of the differentially expressed genes (DEG) of the training cohort were used as input to fit the LASSO Cox regression model with the OS as the response variable. The survival v3.5-3 package was used to create the survival object, which acted as the response variable. Parameter alpha was indicated as 1 to ensure the application of the lasso penalty. The goal of this model was to extract the genes with a prognostic impact that were differentially expressed across MG-stress groups by constructing a penalty function to the residual sum of squares (RSS), which was then multiplied by the regularization parameter lambda (λ). The level of the regularization of the model is controlled by the hyperparameter lambda (λ) value that balances the bias and variance in the resulting coefficients.

Henceforth, coefficient β values were obtained to identify the optimal genes using the 10-fold cross-validation. Based on the risk score of each sample, the cohort was divided into two groups (low-risk with 0–50% vs high-risk with 50–100%). The diagnostic value of these genes was evaluated using receiver operating characteristic (ROC) curves.

### scRNA-seq processing pipeline

Primary tissue datasets (GSE154778, GSE155698, PRJCA001063) were scaled and normalized with SCTransform (Seurat v5.0.1) [28]. The 3,000 most variable features were used for PCA, and the top 30 components for SNN graph construction, clustering, and UMAP embedding. Cohorts were integrated with Harmony (Supplementary Figure 4) [29]. Cluster marker genes were identified with FindAllMarkers and annotated based on the literature.

### Sample processing and transcriptomic profile of ST samples

ST profiling was performed as described by Tourneur et al. [30]. Frozen tumor blocks were sectioned (10 µm) and processed with the spRNA-seq protocol [31]. Sections were mounted on Codelink Activated slides, fixed, H&E-stained, scanned (40×), and permeabilized. After cDNA synthesis, tissue removal, probe release, and library preparation, samples were quality-checked and sequenced on an Illumina NovaSeq 6000. Bright-field H&E images were acquired with a Nanozoomer-SQ scanner (Hamamatsu C13140-01).

### ST-seq processing pipeline

Nanozoomer-SQ images were processed with ndpi2tiff v1.8, tifffastcrop v1.3.10 and vips v8.4.5 to rotate, convert to pyramidal TIFF, and generate JPEGs. ST Spot Detector [32] was used for image alignment and spot detection. A custom affine transform, defined by four reference points on the full-size H&E tissue, interpolated spot coordinates to full-size XY positions. Raw sequencing data was processed with the Salmén et al. pipeline v1.7.6 [33] (alignment, annotation, demultiplexing, filtering) using GRCh38 and Gencode v28. Outputs (UMI counts, corrected spot coordinates, H&E images) were integrated into a Seurat spatial object. Data were normalized with SCTransform (Seurat v5.0.1) and processed following Seurat guidelines. Samples were merged, reduced by PCA, clustered, and annotated using FindAllMarkers. Gene signature scores were calculated with AddModuleScore.

### Functional analysis

Gene Set Enrichment Analysis (GSEA) was performed on a pre-ranked list of genes using the fgsea R package v1.24.0. The enrichment of gene sets with a padj< 0.05 were considered as significant. Gene signatures of PDAC and cancer-associated fibroblasts (CAF) subtypes were collected from the CancerRNASig package. Molecular Signature Database (MSigDB), Ontology and Canonical pathways gene sets were obtained by the msigdb package v1.6.0. Finally, immune cell fractions were estimated by the MCP counter algorithm and statistical analysis between treatments of the immune populations was obtained by the package ggpubr v0.6.0.

### Statistical analysis

Patient characteristics were summarized using descriptive statistics (counts, percentages). Group comparisons were performed using Fisher’s exact or Pearson’s Chi-squared test, depending on expected counts, with gtsummary v1.7.2.

Survival analyses were performed using the survival package (v3.5-3) and survminer package (v0.4.9). Kaplan–Meier curves were compared using log-rank tests (p < 0.05). Multivariate Cox proportional hazards models were used to estimate hazard ratios (95% CI). Overall survival (OS) was defined as the time from diagnosis to death, and disease-free survival (DFS) as the time from diagnosis to recurrence after surgery.Transcriptomic data were analyzed using the Wilcoxon test in R (v4.2.3) and RStudio (v2023.3.0.386), with statistical significance set at p < 0.05. Group comparisons for PAMG and gene expression were performed using the ggpubr package (v0.6.0). [34], [35].

## Results

### Identification of a 16-gene signature and its correlation with clinicopathological findings

Crake et al. [20] MG stress classification was applied to the Puleo cohort [21] (n=309) allowing the classification of PDAC tumors with MG high (n=45) or MG low (n=44) status, as detailed in the materials and methods section. Differential gene expression (DGE) analysis identified 365 differentially expressed genes between the two groups (Figure 1). Impact prognostic genes were then selected using a LASSO Cox regression model based on overall survival (OS) (Supplementary Figure 1-A). The optimal regularization parameter (λ) was chosen by ten-fold cross-validation at the λ minimum error (Supplementary Figure 1-B). Sixteen genes with non-zero coefficients were retained (*FAM162A, SEC61G, NT5E, TREM1, SLC3A1, CALB2, TFF1, SLC11A1, PDIA2, PADI1, ADH1C, FXYD3, SCEL, ECT2, TIPARP, IGHG1*) (Supplementary Table 1).

A risk score was calculated from the expression and coefficients of these genes. Based on the median score, patients were divided into MG-GEM high (n=43) and low (n=42) groups (Figure 2-A). The model showed high predictive accuracy (AUC: 0.854 at 2 years, 0.778 at 4 years, 0.753 at 6 years) (Supplementary Figure 1-C).

**Figure 2.**
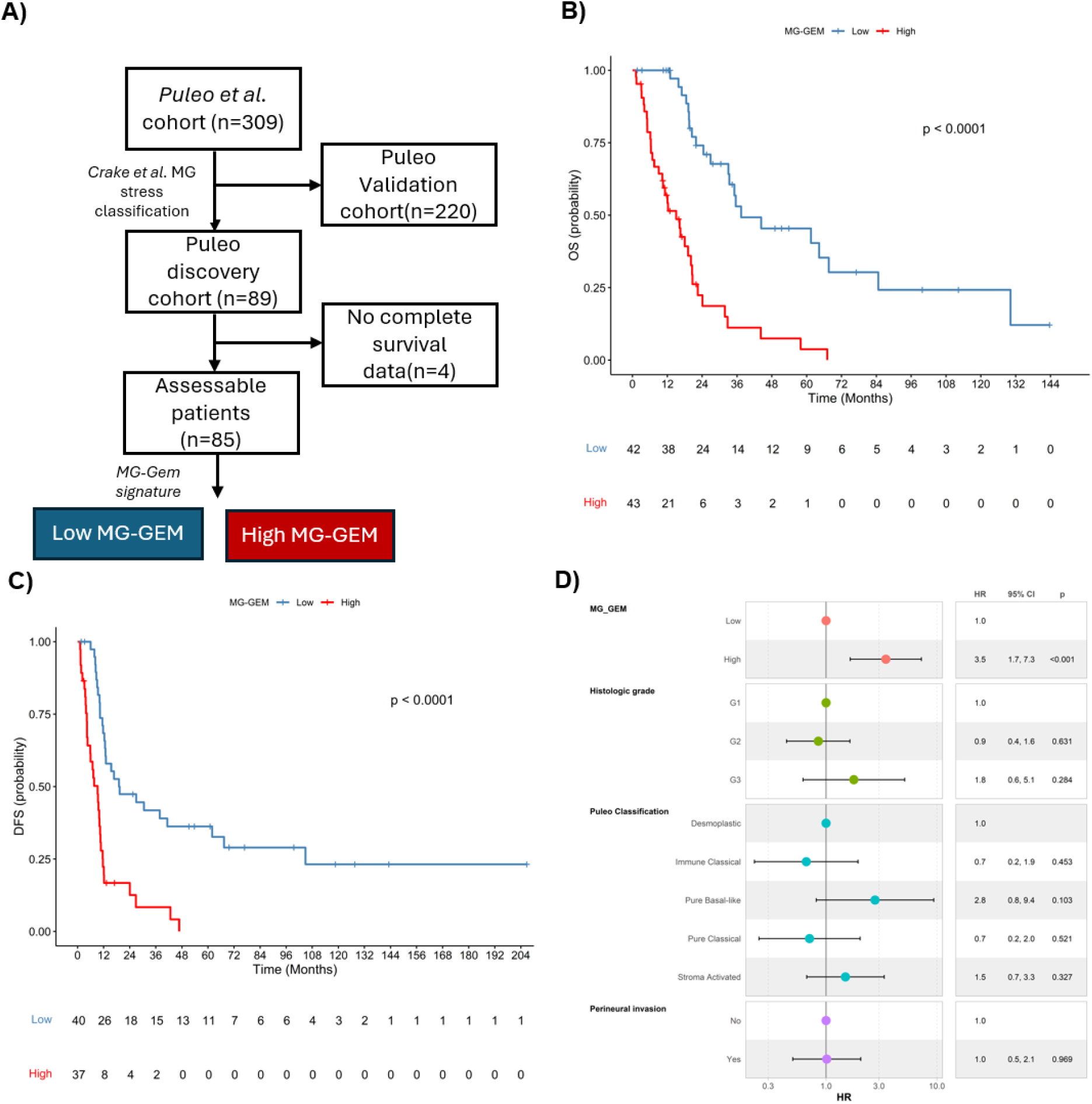
Survival analysis of MG-GEM stratification in the Puleo discovery cohort (n = 89) **(A)** Flowchart of the discovery cohort. **(B, C)** Kaplan-Meier analysis of overall survival (OS) and disease-free survival (DFS). **(D)** Forest plot showing multivariate Cox proportional hazards regression including MG-GEM groups. **Abbreviations:** MG, Methylglyoxal; GEM, Gemcitabine; OS, Overall Survival; DFS, Disease-Free Survival; HR, Hazard Ratio; CI, Confidence Interval.

Univariate analysis of CP findings revealed that MG-GEM status was significantly associated with perineural invasion (high: 85%, low: 66%, p=0.040) (Table 1). In the discovery cohort, high MG-GEM patients had significantly shorter OS (*11.7 vs. 27.2 months, p<0.0001*) and DFS (*7.6 vs. 17.8 months, p<0.0001*) compared with low MG- GEM patients (Figure 2-B,2-C).

**Table 1.** Clinical Characteristics of the high and low MG-GEM patient groups.

| Variable | N | MG-GEM |  |  | p-value<sup>2</sup> |
| --- | --- | --- | --- | --- | --- |
|  |  | Overall N = 85<sup>1</sup> | High Risk N = 43<sup>1</sup> | Low Risk N = 42<sup>1</sup> |  |
| **Age** | 85 |  |  |  | 0.23 |
| <= 65 years |  | 35 (41%) | 15 (35%) | 20 (48%) |  |
| >65 years |  | 50 (59%) | 28 (65%) | 22 (52%) |  |
| **Sex** | 85 |  |  |  | 0.72 |
| Female |  | 34 (40%) | 18 (42%) | 16 (38%) |  |
| Male |  | 51 (60%) | 25 (58%) | 26 (62%) |  |
| **Histological grade** | 83 |  |  |  | 0.091 |
| G1 |  | 27 (33%) | 9 (21%) | 18 (44%) |  |
| G2 |  | 42 (51%) | 25 (60%) | 17 (41%) |  |
| G3 |  | 14 (17%) | 8 (19%) | 6 (15%) |  |
| **Resection Margin** | 85 |  |  |  | 0.28 |
| R0 |  | 64 (75%) | 30 (70%) | 34 (81%) |  |
| R1 |  | 19 (22%) | 11 (26%) | 8 (19%) |  |
| R3 |  | 2 (2.4%) | 2 (4.7%) | 0 (0%) |  |
| **TNM.Stage** | 85 |  |  |  | 0.64 |
| IA |  | 3 (3.5%) | 1 (2.3%) | 2 (4.8%) |  |
| IB |  | 4 (4.7%) | 1 (2.3%) | 3 (7.1%) |  |
| IIA |  | 15 (18%) | 7 (16%) | 8 (19%) |  |
| IIB |  | 63 (74%) | 34 (79%) | 29 (69%) |  |
| **Lymphatic invasion** | 82 | 50 (61%) | 23 (56%) | 27 (66%) | 0.37 |
| **Perineural invasion** | 82 | 62 (76%) | 35 (85%) | 27 (66%) | **0.040** |
| **Vascular invasion** | 82 | 42 (51%) | 20 (49%) | 22 (54%) | 0.66 |
| **Adjuvant Treatment** | 85 |  |  |  | 0.43 |
| 5FU |  | 2 (2.4%) | 1 (2.3%) | 1 (2.4%) |  |
| Gemcitabine |  | 54 (64%) | 24 (56%) | 30 (71%) |  |

| MG-GEM |  |  |  |  |  |
| --- | --- | --- | --- | --- | --- |
| Variable | N | Overall N = 85<sup>1</sup> | High Risk N = 43<sup>1</sup> | Low Risk N = 42<sup>1</sup> | p-value<sup>2</sup> |
| Other |  | 2 (2.4%) | 1 (2.3%) | 1 (2.4%) |  |
| Untreated |  | 27 (32%) | 17 (40%) | 10 (24%) |  |
| <sup>1</sup>Median (IQR) or Frequency (%) |  |  |  |  |  |
| <sup>2</sup>Pearson's Chi-squared test; Fisher's exact test |  |  |  |  |  |

To test whether MG-GEM was an independent prognostic factor, a multivariate Cox model was constructed including histological grade, perineural invasion, and Puleo subtype. MG-GEM remained independently associated with OS (*HR 3.5, 95% CI 1.7– 7.3, p<0.001*) (Figure 2-D).

### Molecular and Transcriptomic Profiling Reveal Distinct Tumor Features Associated with MG-GEM Stratification

To define the molecular landscape of MG-GEM groups, we examined key mutations and transcriptomic subtypes (Supplementary Table 2). *KRAS G12D* was more frequent in high MG-GEM tumors (59% vs. 33%, p=0.043), while *G12V* predominated in low MG-GEM tumors (53% vs. 22% p=0.043) *SMAD4* nonsense mutations were enriched in high MG-GEM (25% vs. 7.9%, p=0.049).

Transcriptomic classifiers consistently linked high MG-GEM tumors to poor-prognosis subtypes. In the Puleo scheme, they were mainly Pure Basal-Like (20% vs 2.7%) and Stroma Activated (38% vs 14%), while low MG-GEM tumors were often Immune Classical (2.5% vs 32%). Similarly, Bailey classification identified most high MG-GEM tumors as squamous (67% vs 9.5%), whereas low MG-GEM tumors were largely progenitor (12% vs 48%) or immunogenic (2.3% vs 14%).

Moffitt stromal profiling showed more activated stroma in high MG-GEM (67% vs. 43%) and normal stroma in low (2.3% vs 19%) (p=0.014). PurIST classification further distinguished the groups: basal tumors were increased in high MG-GEM (33% vs. 8.1%), while classical tumors predominated in low MG-GEM (68% vs. 92%), (p=0.008). GSEA confirmed enrichment of “Basal-like” and “Stroma Active” in high MG-GEM, and “Classical,” “Immune,” and “Stroma Inactive” in low MG-GEM (Figure 3-A).

**Figure 3.**
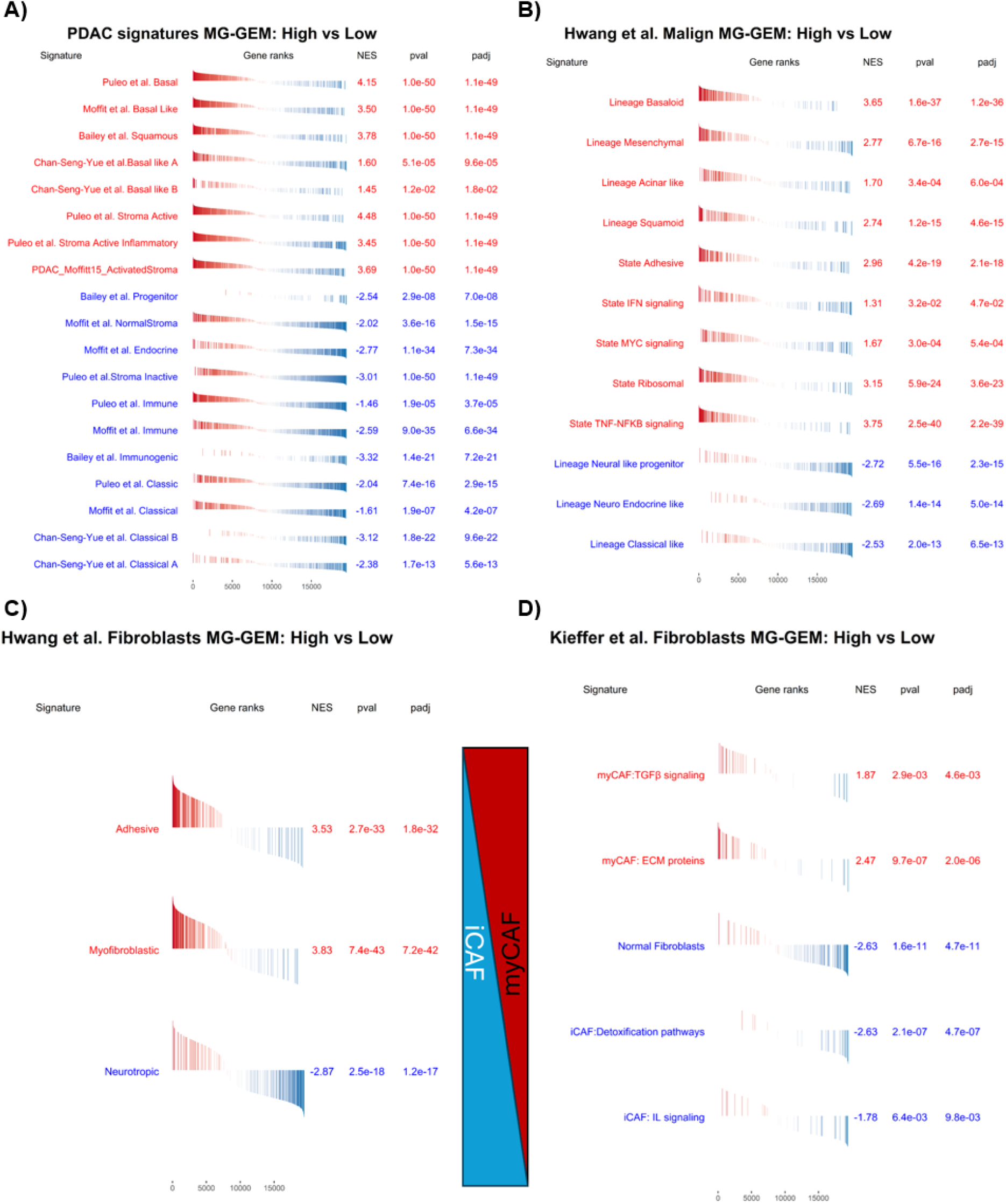
Association of high and low MG-GEM samples with major PDAC molecular subtypes from bulk and single-cell transcriptomic analyses **(A)** Normalized enrichment scores (NES) showing that the high MG-GEM group is significantly enriched for “Basal-like” and “Stroma Active” signatures from bulk molecular classifications. **(B)** NES indicating that the high MG-GEM group is significantly enriched in “Basaloid,” “Mesenchymal,” and “Squamoid” single-cell–derived signatures from Hwang et al. **(C, D)** NES from GSEA of CAF signatures (Hwang et al. and Kieffer et al.) showing enrichment of myCAFs in high MG-GEM samples and enrichment of iCAFs in low MG-GEM samples.

We also explored the utility of MG-GEM in relation to tumor aggressiveness using the Pancreatic Adenocarcinoma Molecular Gradient (PAMG) score [26]. Consistently, PAMG scores were higher in low MG-GEM tumors (p=2.4e-06), reflecting a less aggressive phenotype (Supplementary Figure 1-D). High MG-GEM tumors also showed enrichment of poor-outcome Hwang et al. [36] single-cell signatures (“Basaloid,” “Mesenchymal,” “Squamoid”), while low MG-GEM tumors were enriched in favorable ones (“Neuroendocrine-like,” “Classical-like”) (Figure 3-B).

Next, we further localized MG-GEM gene expression to tumor epithelial cells. In 7 spatial transcriptomics (ST) samples from Erasme repository, MG-GEM genes were highly expressed in basal, acinar, and B cell–associated clusters (Supplementary Figure 2-A) (Supplementary Figure 3). Integration of three scRNA-seq cohorts confirmed expression restricted to tumor epithelial cells (Supplementary Figure 2-B) (Supplementary Figure 4).

### MG GEM is correlated with immune and metabolic contexture in High MG GEM tumors

MCP-counter deconvolution showed a higher abundance of fibroblasts in high MG- GEM tumors (Supplementary Figure 5-A). CAF subtype analysis revealed enrichment of Hwang et al. [41] Adhesive and Myofibroblastic CAFs (myCAFs) signatures in high MG-GEM, while low MG-GEM tumors were enriched in “Neurotropic” CAFs (Figure 3- C). Using Kieffer et al. [37] signatures, GSEA confirmed predominance of myCAF programs in high MG-GEM (“TGFβ signaling,” “ECM proteins”), whereas low MG-GEM samples were enriched in normal fibroblast and inflammatory CAF (iCAF) signatures (Figure 3-D).

Metabolic profiling by Karasinska et al. [15] subtypes classified most high MG-GEM tumors as glycolytic (75% vs. 25%, p=0.004), and low MG-GEM tumors as cholesterogenic (78% vs. 22%, p=0.004) (Supplementary Figure 6-A). MG-GEM expression correlated with glycolytic programs in the high group (R²=0.51, p<0.0001) and with cholesterogenic programs in the low group (R²=0.53, p<0.0001) (Supplementary Figure 6-B). Consistently, “REACTOME_GLYCOLYSIS” was enriched in high MG-GEM tumors (Supplementary Figure 6-C).

Gene Ontology (GO) analysis showed that high MG-GEM tumors were enriched for hypoxia, adhesion, collagen, and apoptosis, while low MG-GEM tumors were enriched for ion channel and lipid antigen pathways (Supplementary Figure 5-B). Reactome confirmed these differences: high MG-GEM tumors were enriched in glucose metabolism, hypoxia, mitochondrial activity, and ROS pathways, whereas low MG- GEM tumors were enriched in lipid homeostasis and channel pathways (Supplementary Figure 7-A), confirming the molecular differences among groups.

Immune-related differences were also analyzed. High MG-GEM tumors showed enrichment in innate immune pathways, while low MG-GEM tumors were enriched in adaptive immune signatures (Supplementary Figure 7-B). MCP-counter deconvolution further revealed reduced immune infiltration in high MG-GEM tumors. Consistently, Myeloid dendritic cells, B cells, T cells (including CD8+), were significantly more abundant in the low MG-GEM group (Supplementary Figure 5-A).

### MG-GEM predicts survival in patients who have received adjuvant GEM

To evaluate the prognostic value of MG-GEM, we tested it across independent cohorts. In the remaining Puleo samples (n=211), originally excluded for being classified as intermediate MG stress, Kaplan–Meier analysis confirmed shorter OS (p<0.001) and DFS (p<0.001) in high MG-GEM tumors (n=106) (Figure 4-A) (Figure 4-C). Validation in the Erasme University Hospital cohort (n=42) composed of a series of GEM AT patients with resected PDAC, showed consistent results, with reduced OS (p=0.0025) and DFS (p=0.05) in high MG-GEM patients (Figure 4-D) (Figure 4-F).

**Figure 4.**
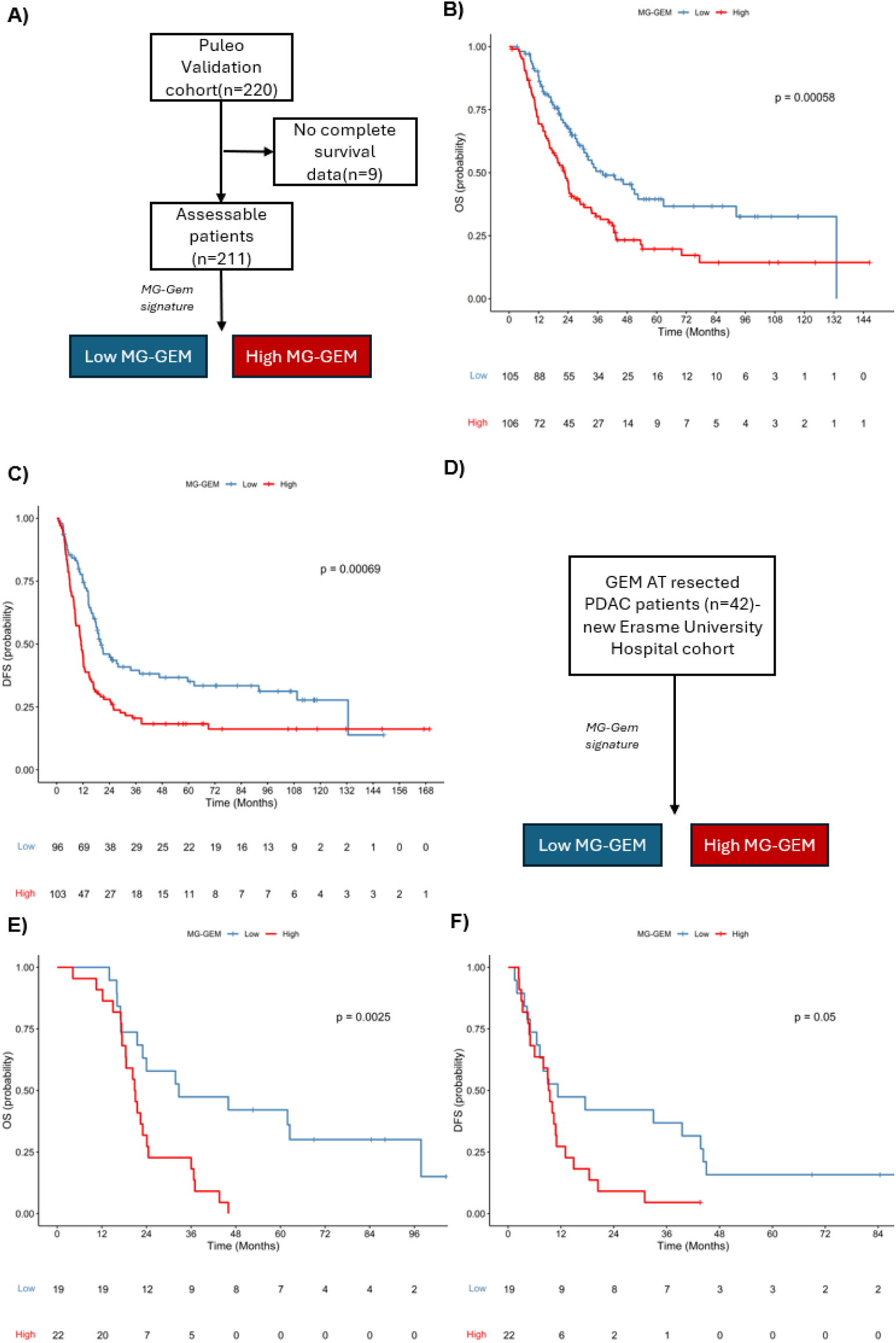
Survival analyses validating the prognostic value of MG-GEM stratification **(A)** Flowchart of the Puleo validation cohort (n = 220). **(B, C)** Kaplan-Meier analysis of the Puleo validation cohort showing significantly worse overall survival (OS). **(D)** Flowchart of GEM-adjuvant resected PDAC patients from the Erasme University Hospital cohort (n = 42). **(E, F)** Kaplan-Meier analysis of the chemotherapy-naive Erasme University Hospital cohort showing significantly worse OS and DFS in the high MG-GEM group. **Abbreviations:** MG, Methylglyoxal; OS, Overall Survival; DFS, Disease-Free Survival.

We next examined the predictive relevance of MG-GEM for GEM response.. Low MG- GEM tumors expressed higher levels of concentrative nucleoside transporter 1 (CNT1) (p=0.0099), a key transporter for GEM uptake, whereas high MG-GEM tumors showed upregulation of ribonucleotide reductase subunit M1 (RRM1) (p=0.047) and cytidine deaminase (CDA) (p=4.9e-05), both associated with GEM resistance (Supplementary Figure 8).

In the Puleo validation cohort, MG-GEM stratified survival only among patients receiving adjuvant GEM (n = 113; Figure 5A), with low MG-GEM patients showing longer overall survival (OS; p = 0.009; Figure 5B) and disease-free survival (DFS; p = 0.012; Figure 5C). No significant differences were observed among patients who did not receive adjuvant therapy (n = 71).Finally, in the PRODIGE-24/TGCC PA6 cohort (183 FOLFIRINOX, 167 GEM) (Figure 5-D), MG-GEM predicted outcome only in GEM-treated patients (OS p=0.0017; DFS p=0.0038) , but not in those treated with FOLFIRINOX (OS p=0.66; DFS p=0.82) (Figure 5-E) (Figure 5-F). These results indicate that MG-GEM is a prognostic biomarker that specifically predicts survival benefit following adjuvant GEM treatment.

**Figure 5.**
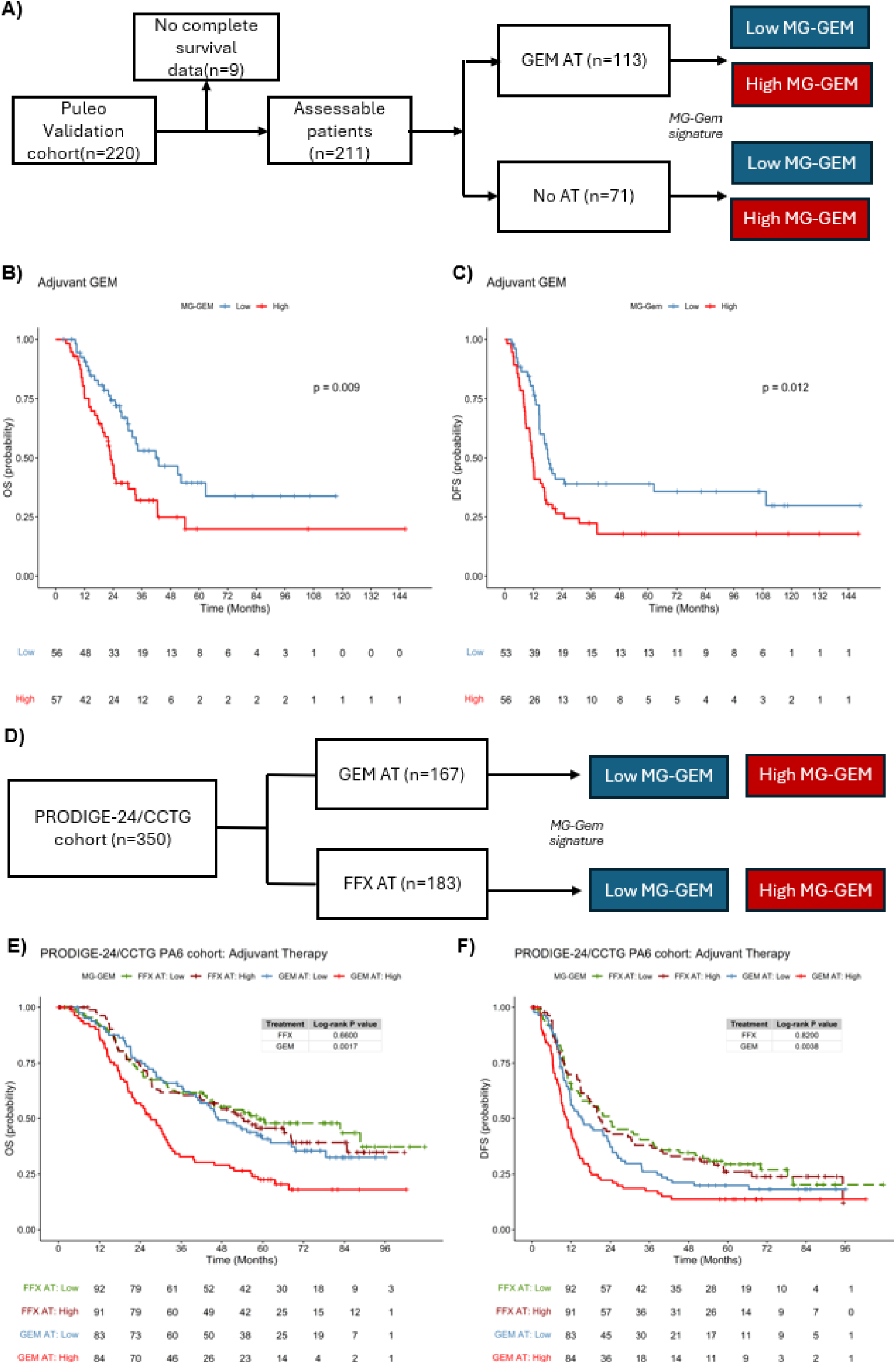
MG-GEM specifically predicts survival benefit from GEM therapy but not to FFX. **(A)** Puleo Validation cohort flowchart stratified by the type of adjuvant treatment. **(B, C)** Kaplan-Meier analysis of the adjuvant gemcitabine–treated Puleo validation cohort showing significantly worse overall survival (OS) and disease-free survival (DFS) in the high MG-GEM group. **(D)** PRODIGE 24/CCTG PA6 cohort flowchart. **(E,F)** Kaplan-Meier analysis of patients from the PRODIGE 24/CCTG PA6 study. Patients stratified as high MG-GEM and receiving adjuvant gemcitabine (red curve) exhibited significantly poorer overall survival (OS) and disease-free survival (DFS) compared to the low MG-GEM group (blue curve). No significant differences were observed among patients who received adjuvant FOLFIRINOX (brown and green curves). **Abbreviations:** MG, Methylglyoxal; OS, Overall Survival; DFS, Disease-Free Survival; AT, Adjuvant Therapy; GEM, Gemcitabine; FFX, FOLFIRINOX.

### Independent and Complementary Predictive Value of MG-GEM and GemPred Signatures

We next compared MG-GEM with the GemPred transcriptomic signature (Figure 6-A) [38]. The two signatures were independent (OR=0.698, p=0.314) (Figure 6-B), allowing their combination. Within GEM AT samples, MG-GEM stratified OS in both GemPred+ (p=0.029) (Figure 6-C) and GemPred− (p=0.041) groups (Figure 6-D). However, no significant DFS differences were observed (Supplementary Figure 10), indicating that the combined prognostic value of MG-GEM and GemPred is stronger for OS than DFS.

**Figure 6.**
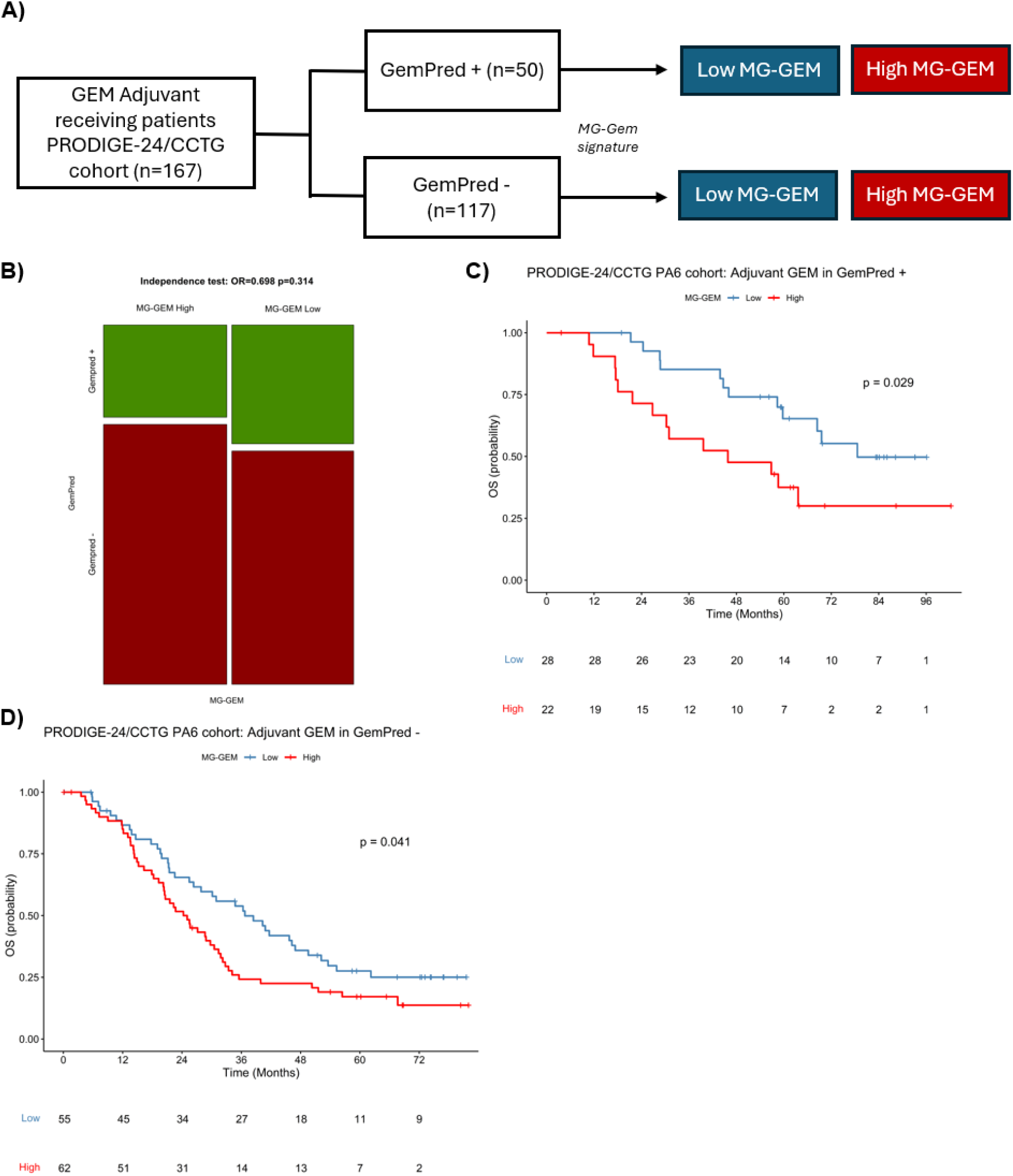
Independent Validation and Integration of MG-GEM and GemPred Signatures (A) Individual flowcharts of the GEM-adjuvant PRODIGE-24/CCTG PA6 cohort stratified by GemPred groups. (B) Fisher’s exact test showing independence between MG-GEM and GemPred groups among samples treated with adjuvant gemcitabine in the PRODIGE-24/CCTG PA6 cohort. (C, D) Kaplan-Meier analysis of the adjuvant gemcitabine–treated PRODIGE- 24/CCTG PA6 cohort showing significantly worse overall survival (OS) in the high MG- GEM group within the GemPred+ (C) and GemPred– (D) subgroups. **Abbreviations**: OS; Overall Survival; MG: Methylglyoxal; AT: Adjuvant therapy; GEM: Gemcitabine.

## Discussion

Several transcriptomic signatures have been developed to predict GEM response in PDAC, including GemPred, GemCore, and PancreasView, each providing complementary insights into tumor biology and chemosensitivity. In this study, we identified differentially expressed genes between PDAC tumors with high and low MG scores that most strongly impacted prognosis, leading to the characterization of a novel 16-gene MG stress signature termed MG-GEM. Given the established connection between MG stress and GEM resistance, we investigated whether MG- GEM signature, could serve as a predictive tool for survival benefit to adjuvant GEM therapy and as a prognostic marker in patients with resected PDAC. Stratification into high and low MG-GEM groups identified patient populations with markedly different prognoses, which was validated across two internal and one external independent cohort, the PRODIGE24/CCTG PA6 trial cohort.

Our MG-GEM signature adds a metabolically informed perspective, capturing glycolytic stress and KRAS-driven rewiring that is not fully represented in these existing tools, and represents, to the best of our knowledge, the shortest transcriptomic signature identified to date for predicting GEM response.. Comparison with the GemPred signature [11], demonstrated that MG-GEM provides independent predictive information. Moreover, combining MG-GEM with GemPred improved the identification of GEM-sensitive and GEM-resistant patients, supporting a complementary role for metabolically informed and clinically derived predictors in personalized treatment strategies. Future clinical translation will require standardized RNA-based assays, consideration of tissue quality and availability, and rapid turnaround to enable timely stratification of patients for adjuvant or neoadjuvant therapies.

Transcriptomic comparison with established PDAC classifications [21], [36], [40], [41], [42], [43], [44], revealed that high MG-GEM tumors were significantly enriched in “Basal-like” and active stroma subtypes, both linked to poor prognosis. In contrast, low MG-GEM tumors were enriched in the “Classical” and immune subtypes, which are associated with better outcomes. Interestingly, a subset of “Pure Classical” tumors fell into the high MG-GEM group, identifying patients who may experience poorer outcomes despite their classical subtype classification. These findings highlight the potential of MG-GEM to refine patient stratification and inform treatment decisions beyond existing classical/basal-like molecular subtyping.

Beyond transcriptomic signatures, additional biomarkers have been explored to guide adjuvant therapy in PDAC. For example, immunohistochemical (IHC) assessment of GATA6, validated in large ESPAC cohorts, has been shown to identify classical versus basal-like subtypes with prognostic and predictive implications [39]. The MG-GEM signature integrates transcriptomic data to reflect functional tumor biology, offering a potential complement to existing IHC and molecular markers.

In PDAC, KRAS mutations contribute to immune evasion and metabolic dysregulation, both of which are key drivers of therapeutic resistance and disease progression [45], [46]. Molecular profiling of MG-GEM-defined subgroups revealed associations with KRAS and SMAD4 mutations. Specifically, KRAS G12D variant was predominantly enriched in the high MG-GEM group, while KRAS G12V was more frequently observed in the low group. Interestingly, a large-scale study investigating the molecular features and clinical outcomes of PDAC patients stratified by KRAS variant revealed that KRAS G12D tumors exhibited a distinct molecular profile, including elevated expression of genes involved in glucose and glutamine metabolism [47].

This suggests that the MG-GEM signature may capture functional heterogeneity downstream of specific KRAS variants. In addition, SMAD4 alterations, which have been implicated in promoting glycolysis [48], and conferring drug resistance through enhanced lipid accumulation [49] were also significantly enriched in the high MG-GEM group. Taken together, these mutational patterns not only provide mechanistic support for the link between MG metabolism and therapeutic outcome but also reinforce the biological plausibility of the MG-GEM signature as a clinically relevant marker of GEM resistance.

Clinically, MG-GEM stratification revealed significant differences in OS and DFS among GEM-treated patients, but not among those treated with FFX, suggesting its specificity for predicting survival benefit from GEM.

Our demonstration of a connection between acquired GEM resistance and MG stress highlighted the involvement of the heat shock response in preclinical PDAC models [20]. Nonetheless, we do not exclude the contribution of additional MG stress-driven mechanisms underlying GEM resistance in glycolytic pancreatic cancer cells. Here, we observed that tumors classified as high MG-GEM exhibited elevated expression of GEM metabolic blockers [20], including CDA and RRM1, compared to low MG-GEM tumors. This expression pattern is consistent with a chemoresistant phenotype and suggests a molecular mechanism linked to the interference with GEM efficacy, warranting further investigation.

This study has limitations, including its reliance on retrospective cohorts and incomplete clinical annotation in some datasets. Future research integrating MG-GEM with comprehensive molecular profiling or predictive biomarkers, as well as evaluating its performance in the neoadjuvant setting, will be important to establish its clinical utility.

In conclusion, we present a novel, biologically grounded MG-GEM signature that captures glycolytic stress–related mechanisms of chemoresistance in PDAC and is associated with prognostic molecular, immune and metabolic features. Validated across independent cohorts, this 16-gene model classifies tumors according to survival benefit from GEM therapy and provides information beyond established molecular subtypes. Importantly, this novel signature warrants further investigation as a potential clinical tool for guiding chemotherapy decisions and opens avenues for exploring KRAS-targeted combination strategies in pancreatic cancer.

## DATA SHARING STATEMENT

The datasets that support the findings of this study are not publicly available. Access to the dataset will be granted upon reasonable request sent to the corresponding author (OAS) and Unicancer.

## AUTHOR CONTRIBUTIONS

Conception and design:

Oier Azurmendi Senar, Akeila Bellahcène, Tatjana Arsenijevic, Jean-Luc Van- Laethem.

Financial support:

Akeila Bellahcène, Jean-Luc Van-Laethem.

Administrative support:

Marjorie Mauduit, Tatjana Arsenijevic.

Provision of study materials or patients:

Christelle Bouchart, Julie Navez, Thierry Conroy, Jérome Cros, Jean-Luc Van- Laethem.

Collection and assembly of data:

Kosta Stosic, Jawad Tarfouss, Laurine Verset, Tatjana Arsenijevic.

Data analysis and interpretation:

Oier Azurmendi Senar, Rémy Nicolle, Vincent Detours, Tatjana Arsenijevic.

Manuscript writing: All authors

Final approval of manuscript: All authors

Accountable for all aspects of the work: All authors

## Acknowledgments

We thank Unicancer and MOSAPAC consortia for providing PRODIGE-24/CCTG PA6 clinical data.

## Supplementary Tables

**Supplementary table 1.**
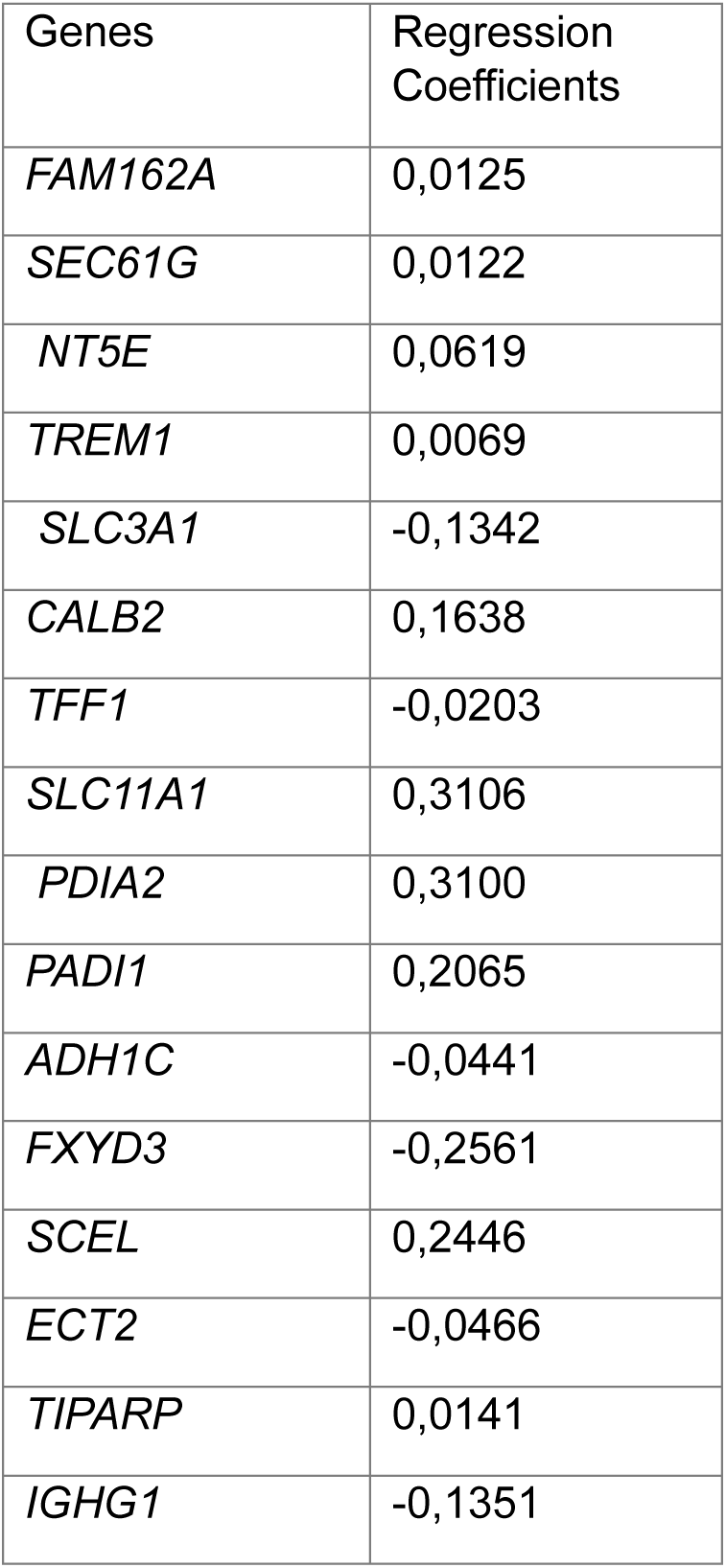
LASSO Cox regression coefficients of the 16 genes used, together with their expression levels, to calculate the final MG-GEM risk score.

**Supplementary Table 2.** Molecular characteristics of patients stratified by high and low MG-GEM groups.

| Variable | N | MG-GEM |  |  | p-value<sup>2</sup> |
| --- | --- | --- | --- | --- | --- |
|  |  | Overall N = 85<sup>1</sup> | High Risk N = 43<sup>1</sup> | Low Risk N = 42<sup>1</sup> |  |
| **KRAS mutation** | 62 |  |  |  | **0.043** |
| G12D |  | 29 (47%) | 19 (59%) | 10 (33%) |  |
| G12V |  | 23 (37%) | 7 (22%) | 16 (53%) |  |
| wt |  | 10 (16%) | 6 (19%) | 4 (13%) |  |
| **TP53 mutation** | 78 |  |  |  | 0.20 |
| nonsense |  | 47 (60%) | 27 (68%) | 20 (53%) |  |
| wt |  | 29 (37%) | 12 (30%) | 17 (45%) |  |
| indel_frameshift |  | 1 (1.3%) | 1 (2.5%) | 0 (0%) |  |
| indel_inframe |  | 1 (1.3%) | 0 (0%) | 1 (2.6%) |  |
| **SMAD4 mutation** | 78 |  |  |  | **0.049** |
| wt |  | 64 (82%) | 29 (73%) | 35 (92%) |  |
| indel_frameshift |  | 0 (0%) | 0 (0%) | 0 (0%) |  |
| nonsense |  | 13 (17%) | 10 (25%) | 3 (7.9%) |  |
| indel_inframe |  | 1 (1.3%) | 1 (2.5%) | 0 (0%) |  |
| **CDKN2A mutation** | 78 |  |  |  | 0.16 |
| wt |  | 63 (81%) | 29 (73%) | 34 (89%) |  |
| indel_frameshift |  | 6 (7.7%) | 5 (13%) | 1 (2.6%) |  |
| nonsense |  | 8 (10%) | 5 (13%) | 3 (7.9%) |  |
| indel_inframe |  | 1 (1.3%) | 1 (2.5%) | 0 (0%) |  |
| **Puleo et al.** | 77 |  |  |  | **<0.001** |
| Desmoplastic |  | 16 (21%) | 10 (25%) | 6 (16%) |  |
| Immune Classical |  | 13 (17%) | 1 (2.5%) | 12 (32%) |  |
| Pure Basal-like |  | 9 (12%) | 8 (20%) | 1 (2.7%) |  |
| Pure Classical |  | 19 (25%) | 6 (15%) | 13 (35%) |  |
| Stroma Activated |  | 20 (26%) | 15 (38%) | 5 (14%) |  |
| **Bailey et al.** | 85 |  |  |  | **<0.001** |

| MG-GEM |  |  |  |  |  |
| --- | --- | --- | --- | --- | --- |
| Variable | N | Overall N = 85<sup>1</sup> | High Risk N = 43<sup>1</sup> | Low Risk N = 42<sup>1</sup> | p-value<sup>2</sup> |
| ADEX |  | 20 (24%) | 8 (19%) | 12 (29%) |  |
| Immunogenic |  | 7 (8.2%) | 1 (2.3%) | 6 (14%) |  |
| Progenitor |  | 25 (29%) | 5 (12%) | 20 (48%) |  |
| Squamous |  | 33 (39%) | 29 (67%) | 4 (9.5%) |  |
| **Moffit et al. Tumor** | 85 |  |  |  | **0.45** |
| Basal |  | 46 (54%) | 25 (58%) | 21 (50%) |  |
| Classical |  | 39 (46%) | 18 (42%) | 21 (50%) |  |
| **Moffit et al. Stroma** | 85 |  |  |  | **0.014** |
| Absent |  | 29 (34%) | 13 (30%) | 16 (38%) |  |
| Activated |  | 47 (55%) | 29 (67%) | 18 (43%) |  |
| Normal |  | 9 (11%) | 1 (2.3%) | 8 (19%) |  |
| **Purist** | 77 |  |  |  | **0.008** |
| Basal |  | 16 (21%) | 13 (33%) | 3 (8.1%) |  |
| Classical |  | 61 (79%) | 27 (68%) | 34 (92%) |  |
| <sup>1</sup>Median (IQR) or Frequency (%) |  |  |  |  |  |
| <sup>2</sup>Fisher's exact test; Pearson's Chi-squared test |  |  |  |  |  |

## Supplementary Figures

**Supplementary Figure 1.**
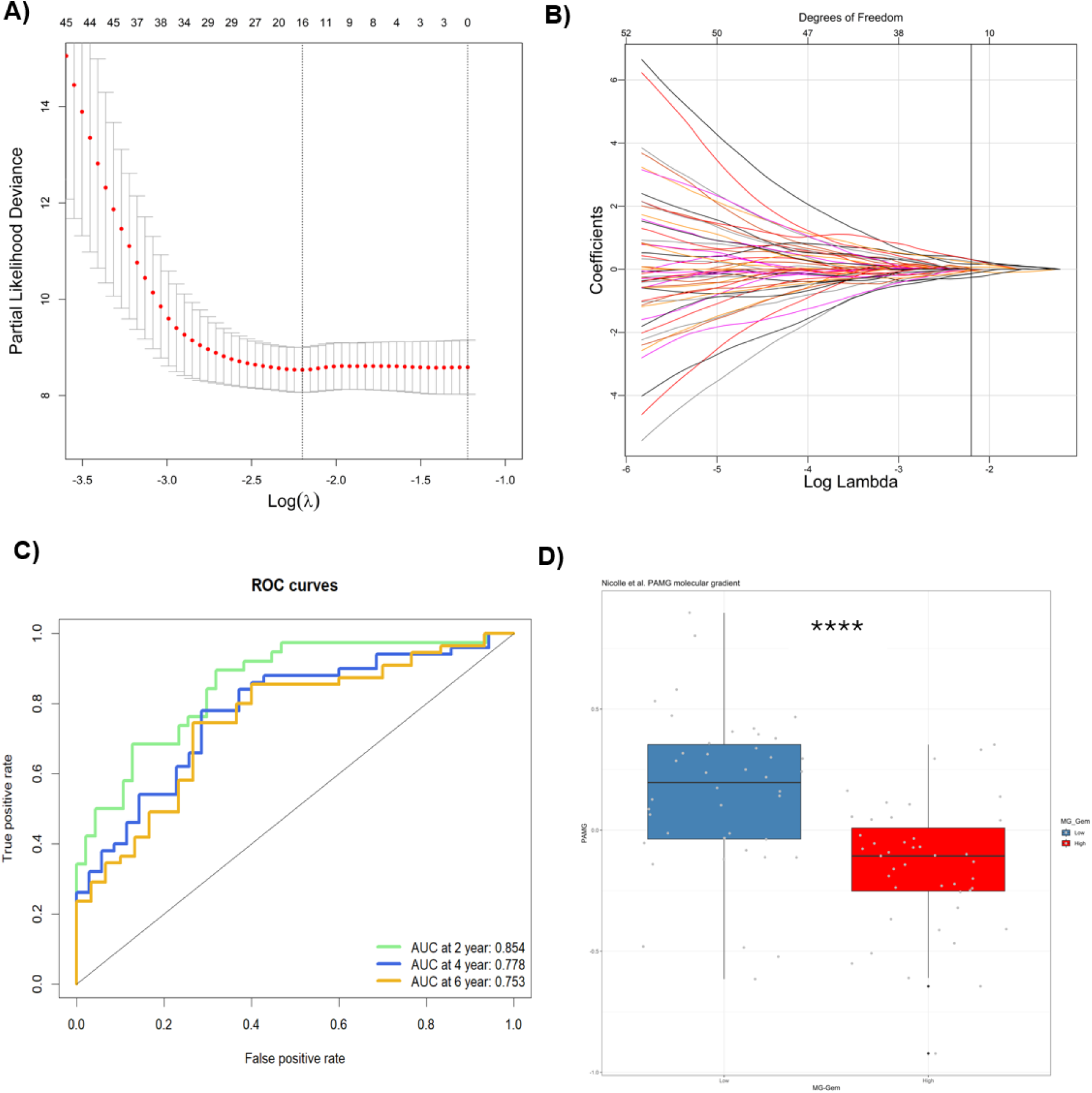
Feature selection and model evaluation using LASSO Cox regression with 10-fold cross-validation (A) LASSO coefficients from the Cox regression model. (B) Coefficient profile plot across the log(λ) sequence. (C) Comparison of ROC curves for the risk score at selected time points. (D) Continuous PDAC molecular gradient (PAMG) applied in the discovery cohort, **showing significantly higher PAMG scores associated with low MG-GEM samples.** Abbreviation: AUC, Area Under the Curve.

**Supplementary Figure 2.**
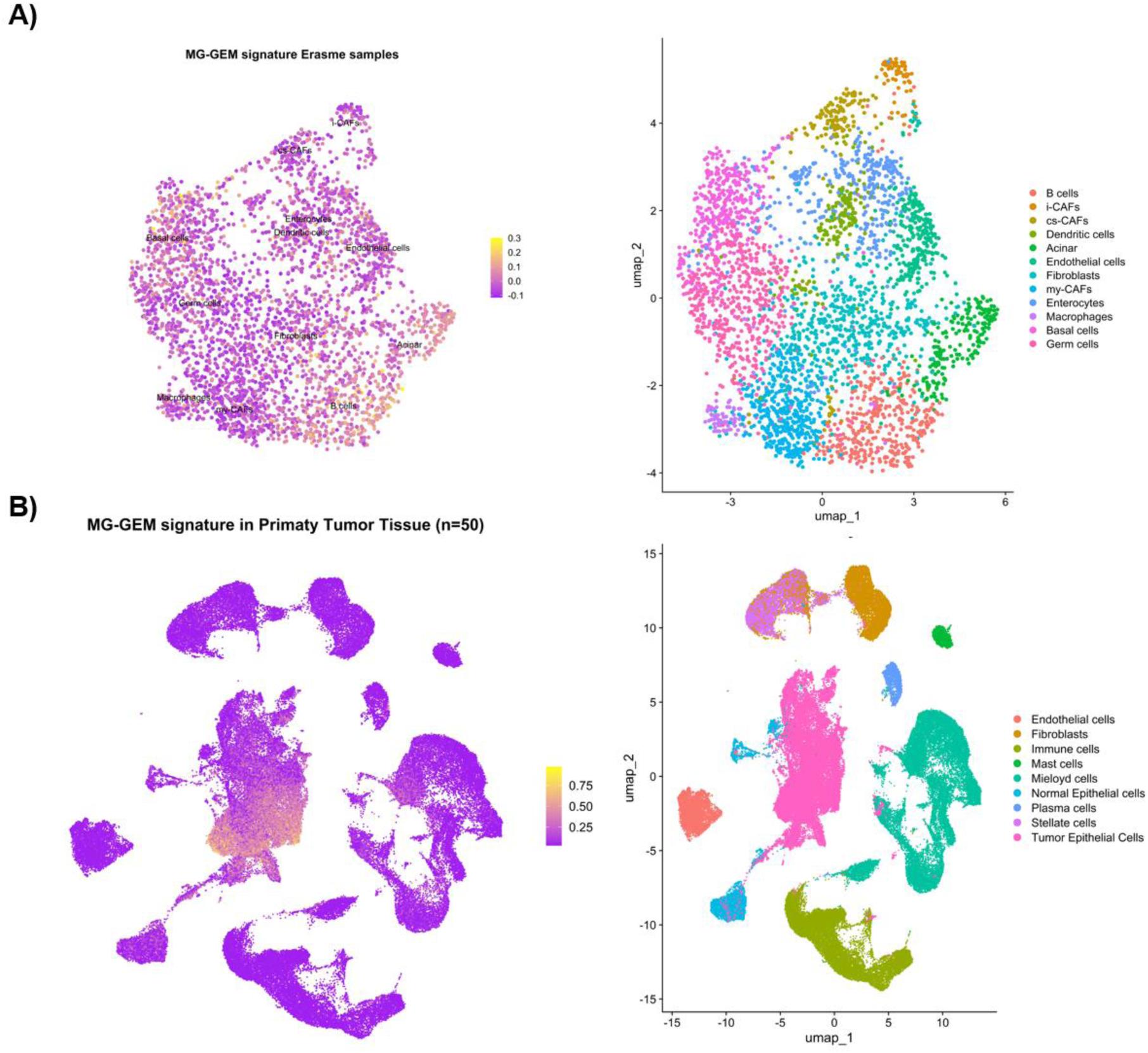
Expression of MG-GEM signature genes in basal cells **(A)** Feature plots of module scores and cluster annotations showing enrichment of MG-GEM signature genes in acinar and basal cells in Erasme ST samples (n = 7). **(B)** Feature plots of module scores and cluster annotations showing enrichment of signature genes in tumor epithelial cells across 50 primary tumor scRNA-seq datasets. **Abbreviations:** MG, Methylglyoxal; ST, Spatial Transcriptomics; scRNA-seq, single-cell RNA sequencing.

**Supplementary Figure 3.**
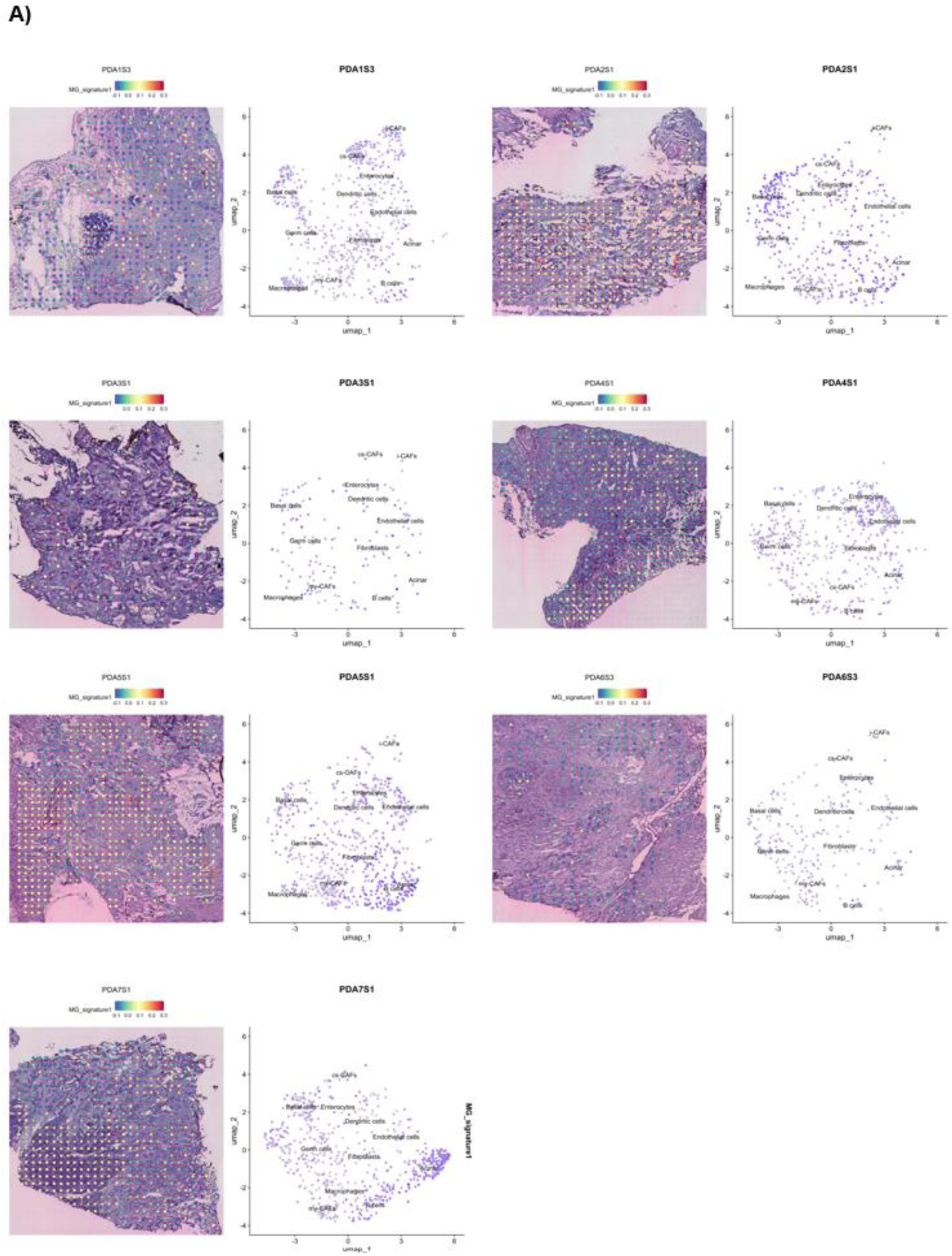
V**a**lidation **of MG-GEM prognostic significance (A)** H&E image and UMAP plots of MG signature gene expression in cohort samples (n = 7).

**Supplementary Figure 4.**
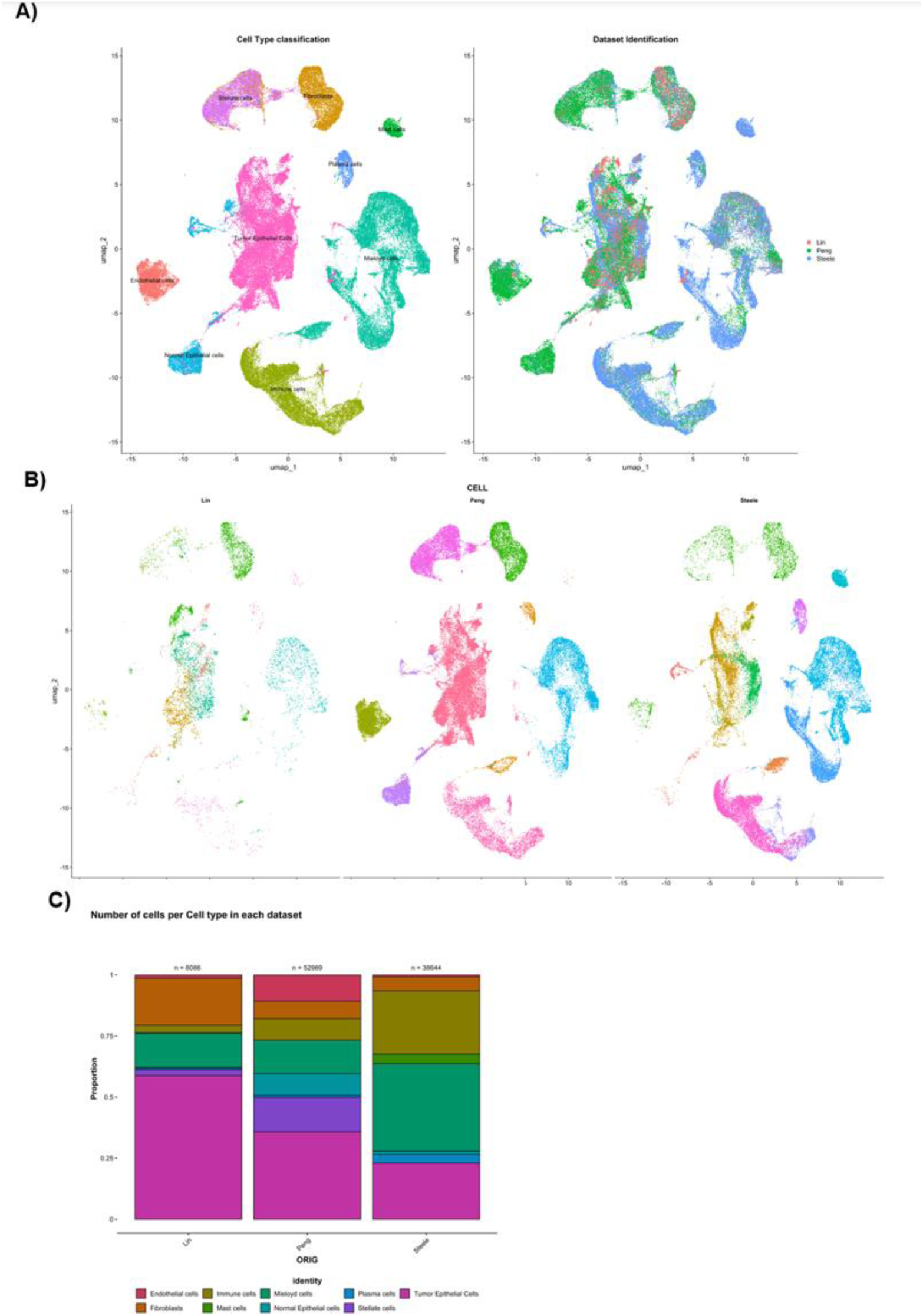
Integration of primary tumor scRNA-seq data from three cohorts **(A)** UMAP plots displaying cell types and original sample IDs in the integrated dataset. **(B)** UMAP plots of each individual dataset showing annotated cell types. **(C)** Bar plots depicting the cell composition of each original dataset. **Abbreviation:** scRNA-seq, single-cell RNA sequencing.

**Supplementary Figure 5.**
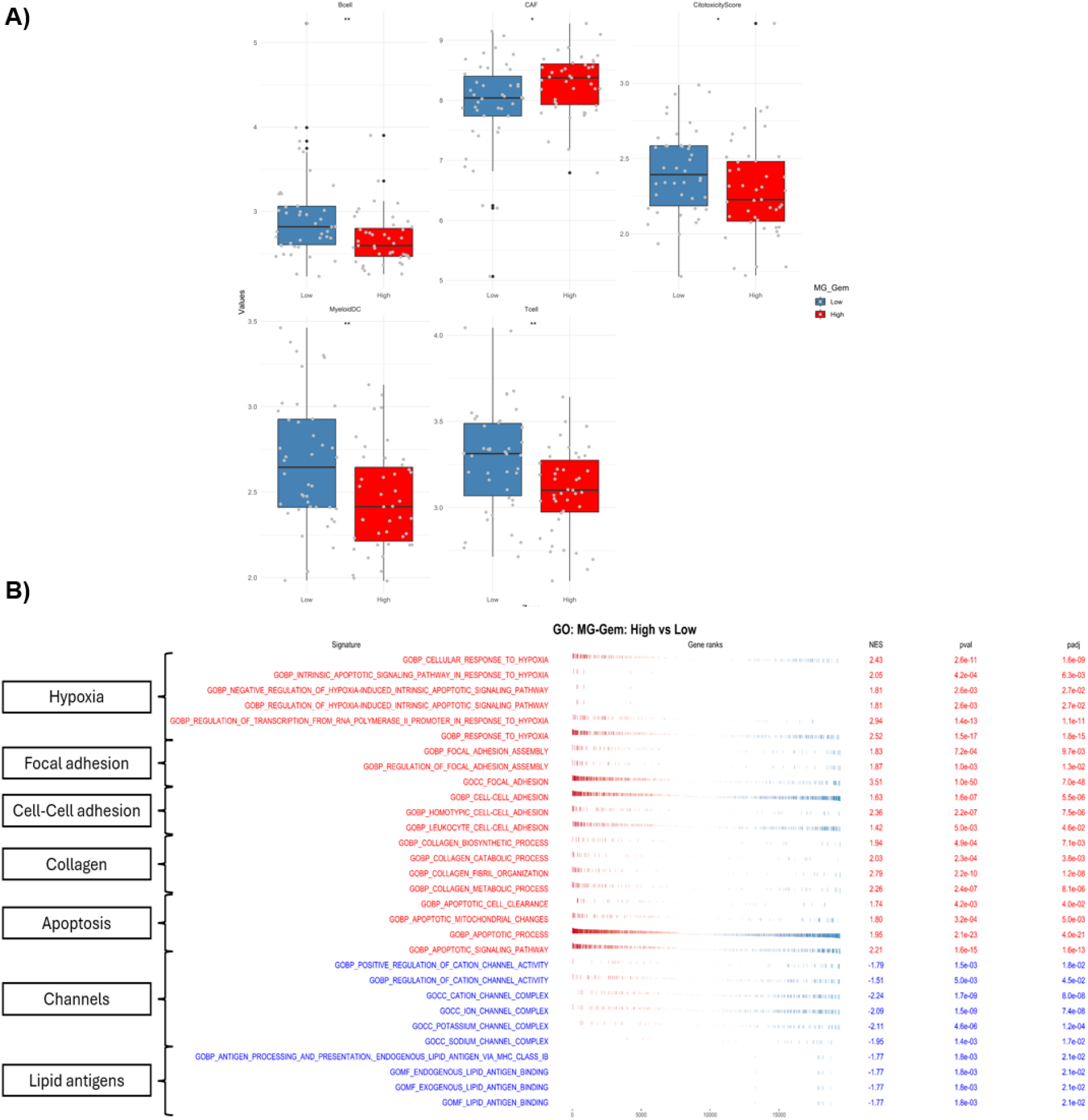
Cancer hallmark signature GSEAs **(A)** Cell type enrichment analysis using MCPcounter revealed significantly lower scores for cytotoxic cells, T cells, B cells, and myeloid dendritic cells, and higher cancer-associated fibroblast (CAF) scores in high MG-GEM samples. **(B)** Normalized enrichment scores (NES) of GO terms related to cancer hallmarks significantly enriched in high MG-GEM versus low MG-GEM samples. **Abbreviations:** MG, Methylglyoxal; GSEA, Gene Set Enrichment Analysis; GO, Gene Ontology.

**Supplementary Figure 6.**
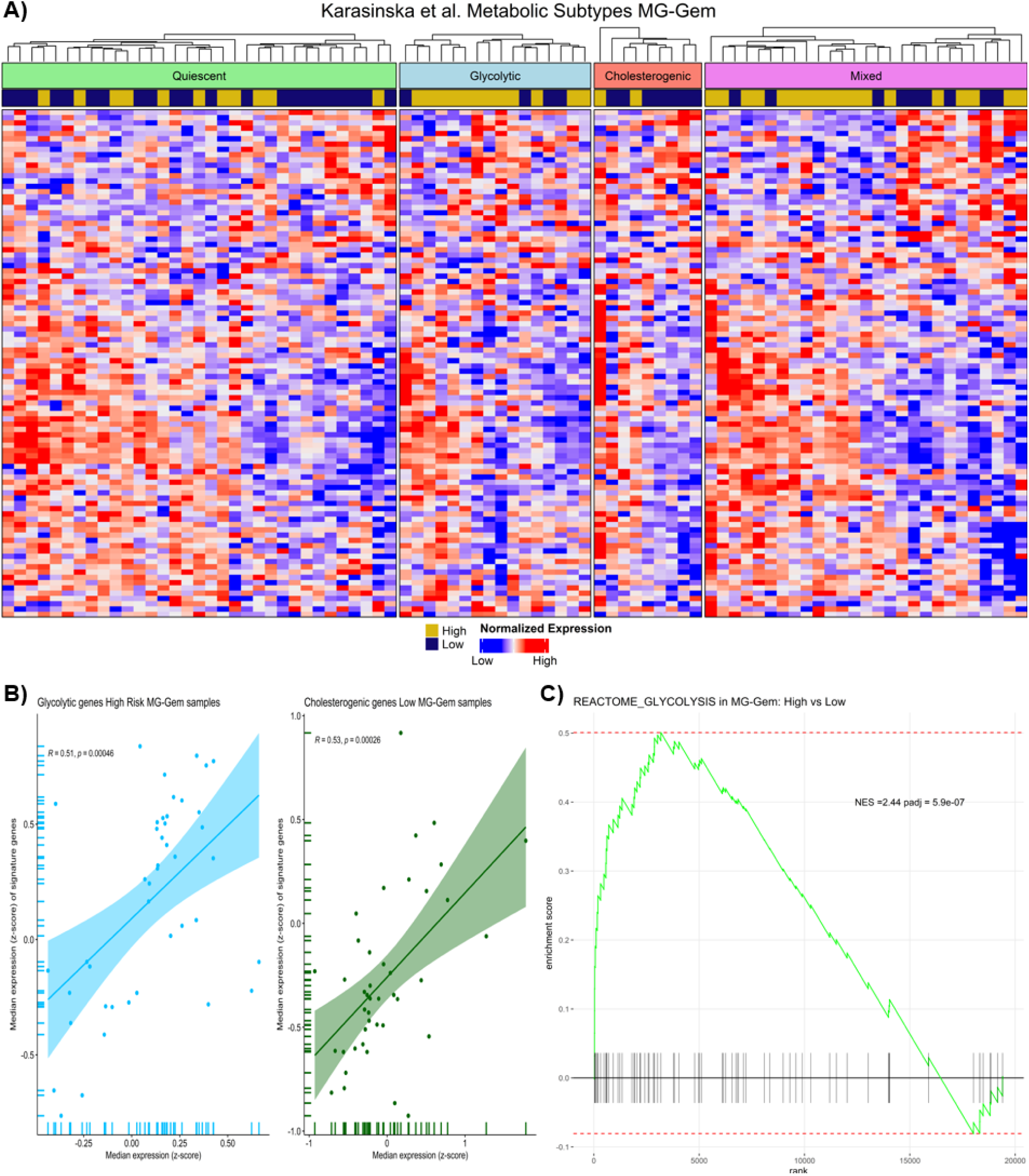
Correlation of MG-GEM groups with Karasinska et al. metabolic subtypes **(A)** Heatmap showing the characterization of Karasinska et al. metabolic subtypes in the Puleo discovery cohort, with glycolytic samples predominantly in the high MG- GEM group and cholesterogenic samples in the low MG-GEM group. **(B)** Positive Pearson correlation between the median expression of glycolytic subtype genes and MG-GEM signature genes in high MG-GEM samples, and a Pearson correlation between cholesterogenic subtype genes and MG-GEM signature genes in low MG-GEM samples. **(C)** GSEA results showing positive enrichment of the pathway **REACTOME_GLYCOLYSIS** in high MG-GEM versus low MG-GEM samples. **Abbreviations:** MG, Methylglyoxal; GSEA, Gene Set Enrichment Analysis.

**Supplementary Figure 7.**
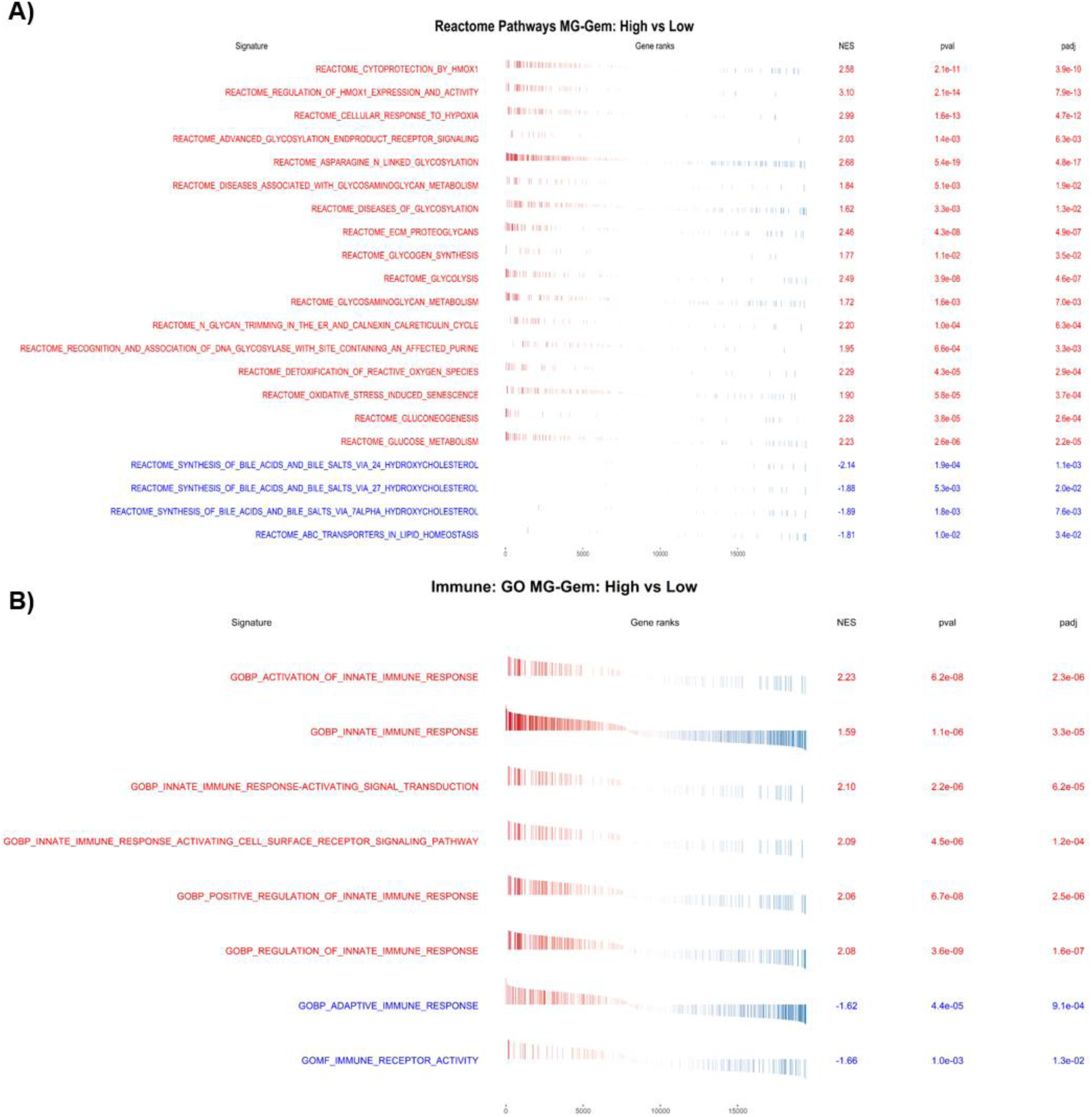
R**e**actome **and immune-related GO term enrichment (GSEA) (A)** Normalized enrichment scores (NES) of Reactome pathways significantly enriched in the high MG-GEM versus low MG-GEM groups. **(B)** NES of immune-related GO terms showing enrichment of innate immunity pathways in the high MG-GEM group and adaptive immunity pathways in the low MG-GEM group. **Abbreviations:** MG, Methylglyoxal; GSEA, Gene Set Enrichment Analysis; GO, Gene Ontology.

**Supplementary Figure 8.**
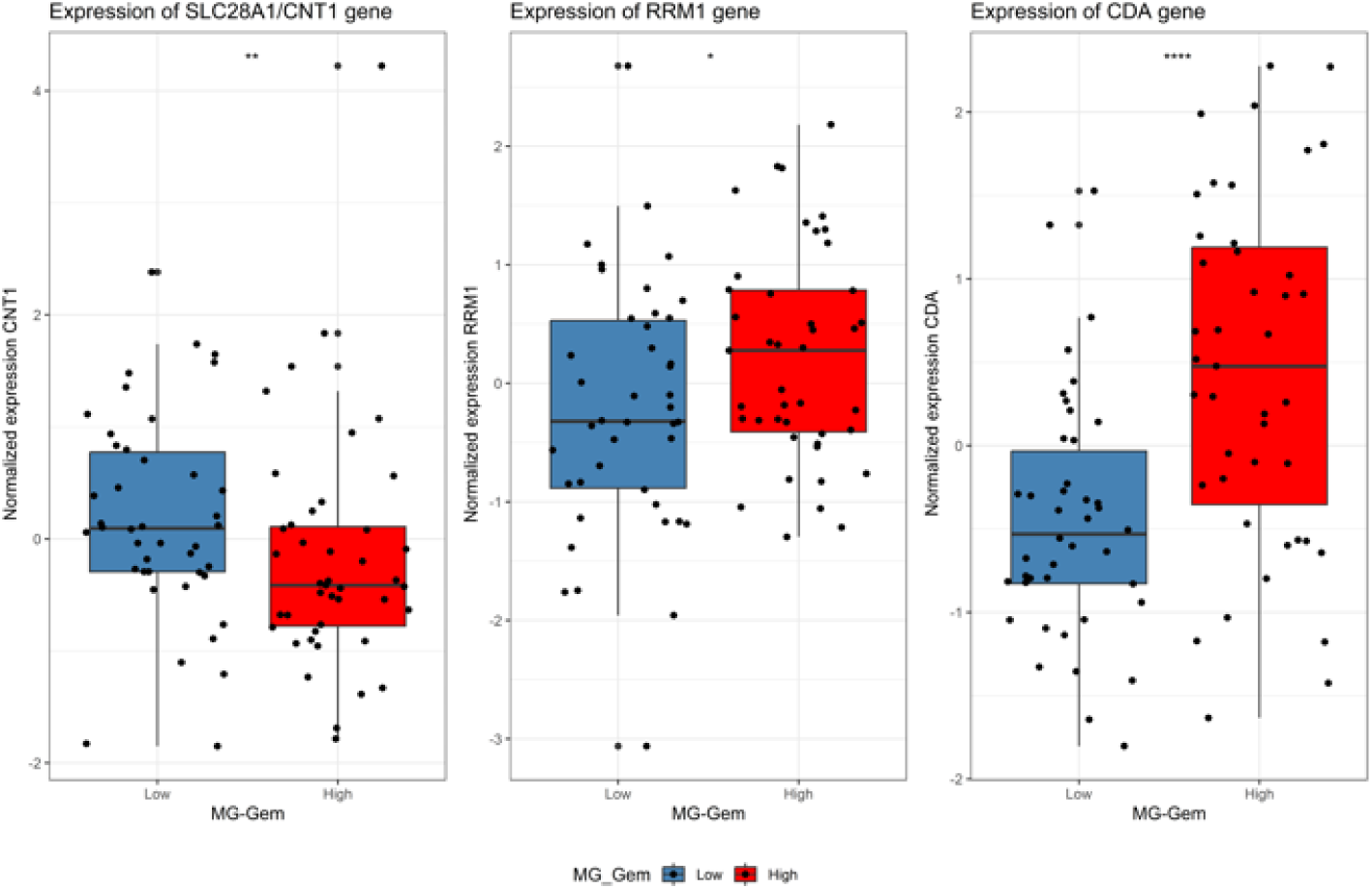
Expression of genes involved in gemcitabine (GEM) transport and metabolism in high and low MG-GEM groups The low MG-GEM group showed higher expression of the GEM transporter CNT1 (p = 0.0099), which mediates GEM uptake. In contrast, the high MG-GEM group exhibited elevated expression of two enzymes associated with GEM resistance: RRM1 (p = 0.047) and CDA (p = 4.9 × 10⁻⁵). Abbreviations: CNT1, Concentrative Nucleoside Transporter 1; RRM1, Ribonucleotide Reductase Catalytic Subunit M1; CDA, Cytidine Deaminase.

**Supplementary Figure 9.**
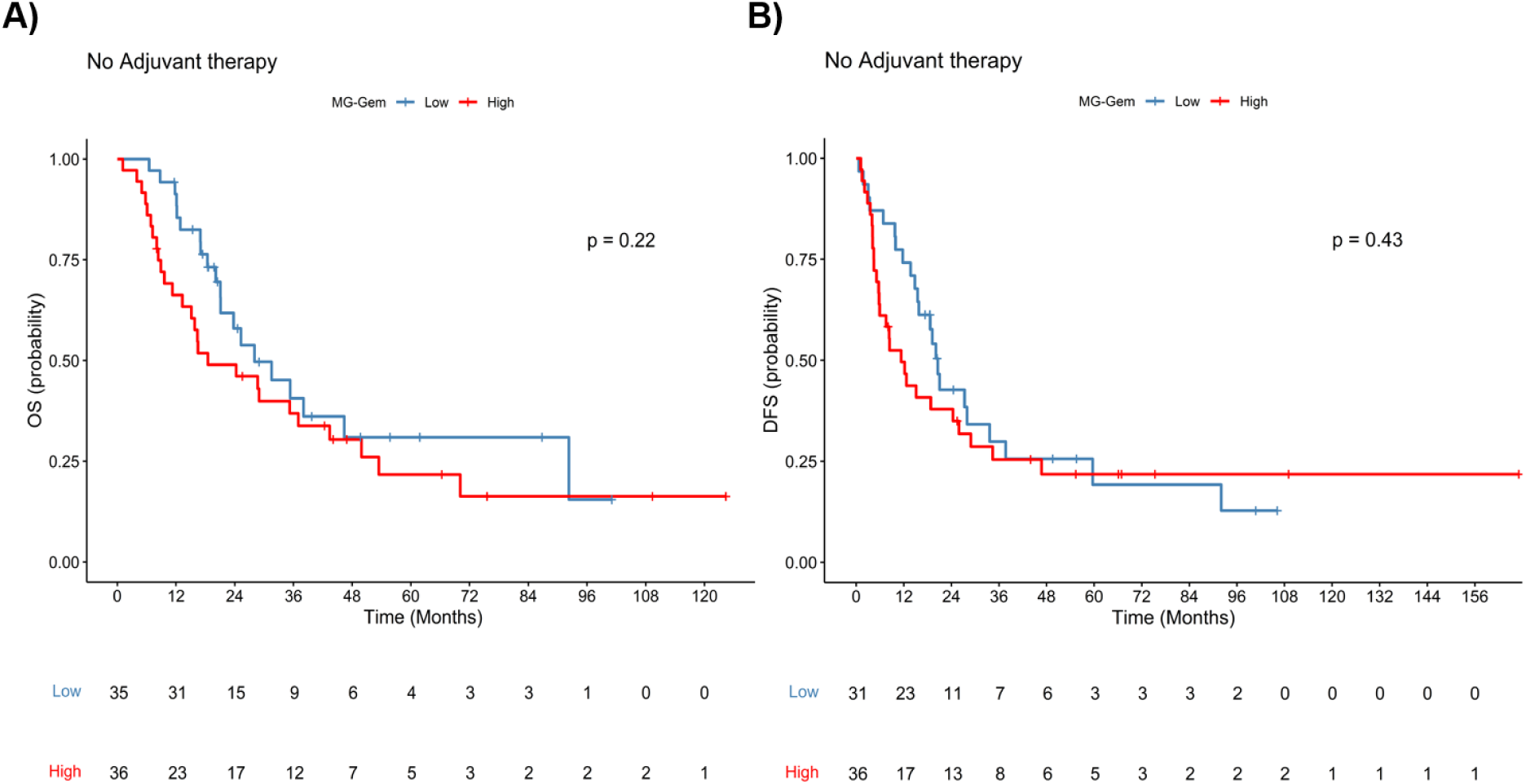
OS and DFS results of the combined MG-GEM and GemPred transcriptomic signatures (A, B) Kaplan-Meier analysis of patients without adjuvant treatment in the Puleo validation cohort showing no significant differences in overall survival (OS, A) or disease-free survival (DFS, B) between high and low MG-GEM groups. **Abbreviations:** MG, Methylglyoxal; DFS, Disease-Free Survival; AT, Adjuvant Therapy; GEM, Gemcitabine.

**Supplementary Figure 10.**
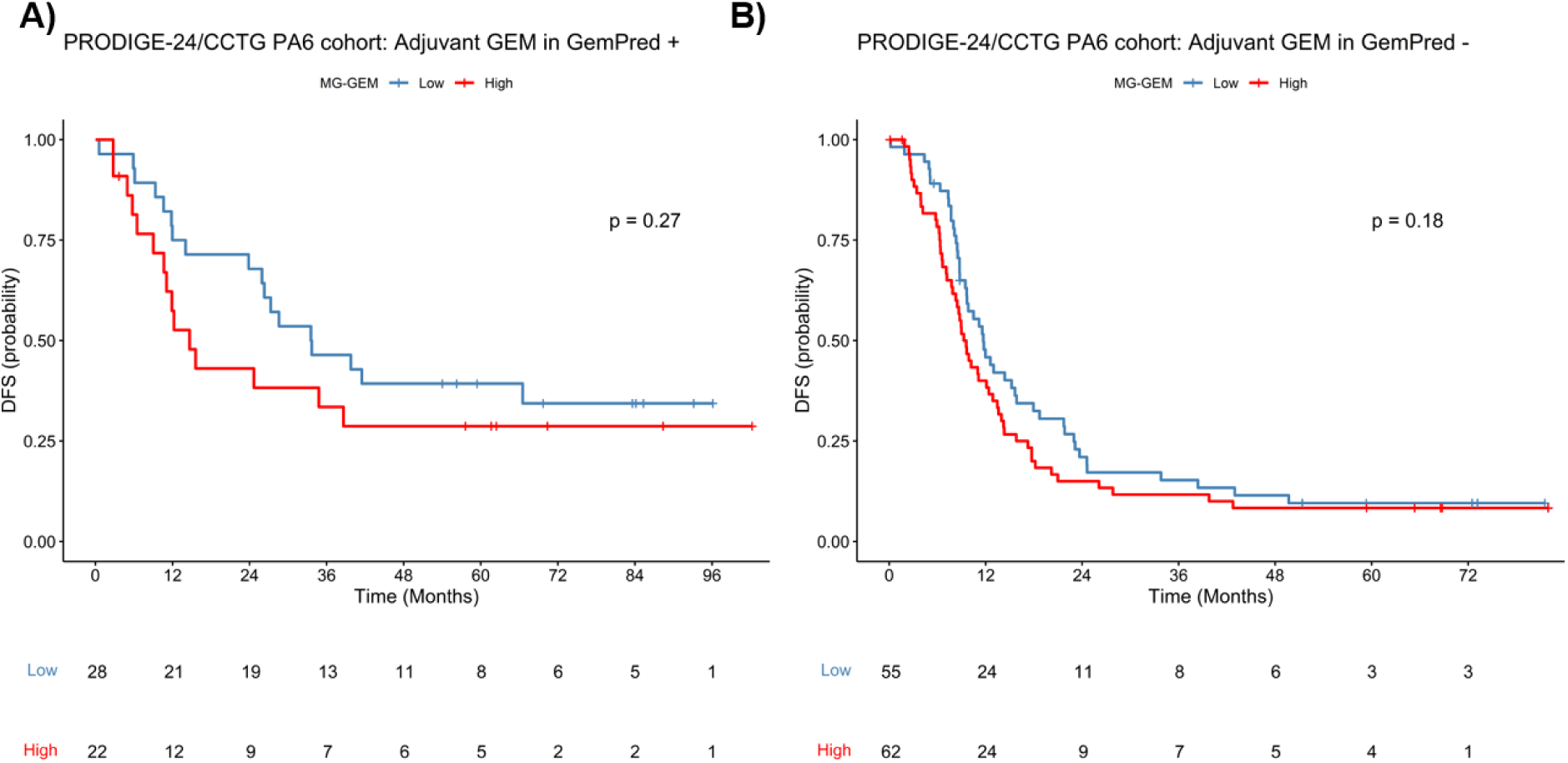
DFS results of the combined MG-GEM and GemPred transcriptomic signatures (A, B) Kaplan-Meier analysis of the adjuvant gemcitabine–treated PRODIGE- 24/CCTG cohort showing no significant differences in disease-free survival (DFS) between high MG-GEM groups within the GemPred+ (A) and GemPred– (B) subgroups. **Abbreviations:** DFS, Disease-Free Survival; MG, Methylglyoxal; AT, Adjuvant Therapy; GEM, Gemcitabine.

